# Connectome-based predictive modeling of cognitive reserve using task-based functional connectivity

**DOI:** 10.1101/2022.06.01.494342

**Authors:** Rory Boyle, Michael Connaughton, Eimear McGlinchey, Silvin P. Knight, Céline De Looze, Daniel Carey, Yaakov Stern, Ian H. Robertson, Rose Anne Kenny, Robert Whelan

**Affiliations:** Department of Neurology, Massachusetts General Hospital, Harvard Medical School; Trinity College Institute of Neuroscience, Trinity College Dublin; Department of Psychiatry, School of Medicine, Trinity College Dublin; School of Nursing and Midwifery, Trinity College Dublin; Global Brain Health Institute, Trinity College Dublin; The Irish Longitudinal Study on Aging (TILDA), School of Medicine, Trinity College Dublin; Cognitive Neuroscience Division, Department of Neurology, Columbia University; Mercer’s Institute for Successful Ageing, St. James’s Hospital

## Abstract

Cognitive reserve supports cognitive function in the presence of pathology or atrophy. Functional neuroimaging may enable direct and accurate measurement of cognitive reserve which could have considerable clinical potential. The present study aimed to develop and validate a measure of cognitive reserve using task-based fMRI data that could then be applied to independent resting-state data. Connectome-based predictive modeling with leave-one-out cross-validation was applied to predict a residual measure of cognitive reserve using task-based functional connectivity from the Cognitive Reserve/Reference Ability Neural Network studies (n = 220, mean age = 51.91 years, SD = 17.04 years). Three network-strength predicted cognitive reserve measures were generated that accurately predicted the residual measures of unseen participants. The theoretical validity of these measures was established via a positive correlation with a socio-behavioural proxy of cognitive reserve (verbal intelligence) and a positive correlation with global cognition, independent of brain structure. This fitted model was then applied to external test data: resting-state functional connectivity data from The Irish Longitudinal Study on Ageing (TILDA, n = 294, mean age = 68.3 years, SD = 7.18 years). The network-strength predicted measures were not positively associated with a residual measure of cognitive reserve nor with measures of verbal intelligence and global cognition. The present study demonstrated that task-based functional connectivity data can be used to generate theoretically valid measures of cognitive reserve. Further work is needed to establish if, and how, measures of cognitive reserve derived from task-based functional connectivity can be applied to independent resting-state data.

## Introduction

Cognitive reserve (CR) refers to a property of the brain that enables better-than- expected cognitive function given the degree of age-related brain changes and brain injury or disease (Collaboratory on Research Definitions for Reserve and Resilience in Cognitive Aging and Dementia, 2022). Higher CR is associated with a delayed onset, and lower incidence, of dementia (Reed et al., 2010; Soldan et al., 2020; Zahodne et al., 2015), and reduced hospitalization risk in individuals with a genetic risk for Alzheimer’s disease (Filshtein et al., 2019). CR is a modifiable construct that may be influenced by various life experiences, for example educational attainment (Malek-Ahmadi et al., 2017) and occupational complexity (Boots et al., 2015), as well as genetics (Barker et al., 2021; Dumitrescu et al., 2020).

Accurate measurement of CR could improve the clinical diagnosis of dementia (Stern, 2012), the measurement of intervention efficacy in clinical trials (Mondini et al., 2016), the stratification of participants in intervention studies (Stern, 2012), and the development of interventions designed to enhance CR (Moga et al., 2019). An accurate neuroimaging measure of CR might enable the identification of specific CR-related brain networks that could be targeted using neuromodulation (Arvaneh et al., 2018; Scheinost et al., 2020) or neurostimulation techniques (Kim et al., 2019).

CR is typically measured using socio-behavioural variables (‘proxies’) that reflect the degree of exposure to various lifetime experiences thought to contribute to CR (Stern et al., 2020). Although this measurement approach is convenient and inexpensive, it is theoretically and methodologically limited as proxies are static self-report variables that fail to capture the entirety of dynamic CR construct (Bettcher et al., 2019; Jones et al., 2011; Ward et al., 2015). Another measurement approach, using structural neuroimaging, is the CR residual, which operationally defines CR as the unexplained variance in cognition after accounting for brain structure and demographics (Bettcher et al., 2019; Reed et al., 2010; Zahodne et al., 2013). In comparison to socio-behavioural proxies, the CR residual can better reflect change in CR over time (Stern et al., 2020). However, the CR residual provides limited insights into the functional processes underlying CR, as it uses structural – not functional – neuroimaging data. Furthermore, because it is a residual, it will necessarily contain a significant proportion of measurement error (Ewers, 2020).

Functional neuroimaging may provide a more direct measure of CR via the identification of neural networks or patterns of neural activity, whose strength or expression differs as a function of CR (Stern et al., 2020; Stern & Barulli, 2019). Unlike proxies, a functional neuroimaging measure could reflect exposure to various lifetime experiences without directly reflecting the change in exposure itself (Stern & Barulli, 2019). This would enable effective evaluation of interventions designed to increase CR. Importantly, a brain-based approach could provide important mechanistic insights into CR.

Valid measurement of CR requires some proposed measure of CR (e.g., a proxy or candidate neuroimaging measure) and two other components: a measure of brain structure/pathology and a measure of cognitive function (Christensen et al., 2008; Stern et al., 2020). The latter two components allow the CR measure to be validated by assessing its protective effect on cognition. A protective effect can be demonstrated by a moderation effect of the candidate CR measure on the relationship between brain structure/pathology and cognition, such that there is a weaker relationship between brain structure/pathology and cognition at higher levels of CR (Collaboratory on Research Definitions for Reserve and Resilience in Cognitive Aging and Dementia, 2022). Alternatively, weaker evidence of a protective effect may be established by a positive association between the candidate CR measure and cognition, controlling for the effect of brain structure/pathology (Stern et al., 2020). Face validity of the CR measure can be established by a positive association with a socio-behavioural CR proxy (Franzmeier, Duering, et al., 2017; Stern & Habeck, 2018).

The theoretical criteria for neuroimaging measures of CR have been satisfied in previous work. Belleville et al. (2021) identified a pattern of increased activation in the right inferior temporal gyrus that was positively associated with a composite CR proxy and that moderated the relationship between hippocampal volume and associative memory performance, i.e. compensating for reduced hippocampal volume. This supports the use of task-fMRI for measuring CR. However, as individual differences in cognitive function are more accurately predicted by global patterns of task-related activations than by regional patterns (Zhao et al., 2021), focusing on globally distributed activations may be a more promising approach. Global brain activation may also be better suited to detecting the generalized neural networks (i.e. generic or task-invariant networks) that may underlie CR (Steffener et al., 2011; Steffener & Stern, 2012; Stern et al., 2018; van Loenhoud et al., 2020).

Resting-state fMRI (rs-fMRI) provides a means of measuring global patterns of connectivity in generic or task-invariant CR networks. In contrast to task-fMRI, rs-fMRI is unaffected by various individual-level factors, including task difficulty (Stern, 2005), motivation, concentration, and fatigue (McCaffrey & Westervelt, 1995), that influence task performance and engagement and therefore may affect task-related activations. As rs-fMRI does not place task-related demands on participants, it can be more easily conducted in individuals with cognitive impairment (Fox & Greicius, 2010), and therefore has better potential for clinical utility. Finally, rs-fMRI data can be more easily shared and aggregated with data from other sites as part of data-sharing initiatives, thereby enabling greater use of any derived CR measures (Mennes et al., 2013; Woodward & Cascio, 2015).

Various studies have identified associations between resting-state connectivity in specific networks and CR proxies as well as protective effects on cognition. Educational attainment has been positively associated with connectivity of the frontoparietal network (FPN; Franzmeier, Caballero, et al., 2017; Serra et al., 2016) and between a salience network (SAL) node, the anterior cingulate cortex, and regions including the right hippocampus, right posterior cingulate cortex/gyrus, left inferior frontal lobe, and left angular gyrus (Arenaza-Urquijo et al., 2013). Greater connectivity of the FPN, SAL and default mode networks (DMN) has been associated with lower cognitive decline, independent of brain structure, and in the context of high amyloid burden (Buckley et al., 2017a).

Converging evidence of the association between resting-state functional connectivity and CR suggests that rs-fMRI may be a viable method for measuring CR. The viability of this method was firmly supported by Stern et al. (2021), who identified a pattern of resting-state functional connectivity that was positively associated with a CR proxy, verbal intelligence. Having demonstrated the face validity of this potential measure, a protective effect on cognition was subsequently established as this measure was associated with global cognition, controlling for cortical thickness. Importantly, Stern et al. were able to validate this measure in an independent dataset, where it also showed face validity and a protective effect on global cognition.

Despite the advantages and demonstrated viability of rs-fMRI data for measuring CR, task-fMRI data may still enable more accurate measurement. Task-based fMRI can augment individual differences in neural processes or networks underlying a phenotype (Greene et al., 2018; Yoo et al., 2018) and has been shown to generate more accurate predictions of cognitive phenotypes compared to rs-fMRI (Greene et al., 2018). Therefore, current methods for developing functional neuroimaging measures of CR may have suboptimal accuracy if developed using rs-fMRI but may have limited clinical potential and shared use if developed using task-fMRI. A novel approach is to develop a measure of CR leveraging the increased accuracy of task-fMRI but that can be applied to rs-fMRI in independent datasets or individual scans, thereby maximizing the clinical potential and usability of the measure.

Connectome-based predictive modeling (CPM; Shen et al., 2017) is a data driven- method for developing accurate measures of cognitive phenotypes using task-based fMRI (or rs-fMRI) data that can be subsequently applied to rs-fMRI data from independent datasets (Rosenberg et al., 2016). In short, CPM summarises the most relevant connections – or ‘edges’ – for the phenotype, across the whole brain. Within cross- validation frameworks, these edges are summed to create three single scalar value measures – positive, negative, and combined network strength – which summarise the connectivity strength of edges that are related to the phenotype of interest. CPM has been widely shown to create measures of cognitive and behavioural phenotypes that generalise across datasets (M. Gao et al., 2020; Rosenberg et al., 2016; Yip et al., 2019; Yoo et al., 2018).

Previous applications of CPM have successfully predicted cognitive phenotypes – fluid intelligence (S. Gao et al., 2019; Greene et al., 2018), attention (Fountain-Zaragoza et al., 2019; Rosenberg et al., 2016), and executive function (Henneghan et al., 2020) – that have been directly associated with CR proxies elsewhere (Chan et al., 2018; Lavrencic et al., 2018). CPM could capitalise on recent developments in measuring CR from neuroimaging data by using the CR residual as the outcome – or target – variable to be predicted from the functional connectivity data. The CR residual has face validity (Habeck et al., 2017; Lee et al., 2019), satisfies the cognitive benefit criterion (Reed et al., 2010; Zahodne et al., 2013) and provides a more direct measure of CR than proxies that have been used as target variables in previous attempts to measure CR with fMRI (Stern et al., 2018, 2021; van Loenhoud et al., 2020). The present study aimed to develop and validate a functional neuroimaging measure of CR by applying CPM to task-based fMRI data to predict a CR residual and to externally validate the measure on resting-state fMRI data in an independent dataset.

## Methods

### Participants

#### Training Set

The training set consisted of data from 220 participants of the Cognitive Reserve/Reference Ability Neural Network (CR/RANN) studies (Stern et al., 2014, 2018). From an initial 384 participants, 123 were excluded due to missing data, presence of possible lesions, or fMRI data quality issues (*see Supplemental Information, Methods: Participant exclusions*) and 41 were excluded for excessive head motion during fMRI scan, defined as mean framewise displacement (FWD) > 0.4 mm or frame to frame movements > 97.5^th^ percentile of frame to frame movements across the whole sample.

#### Test Set

The test set consisted of 294 participants from the MRI subset of The Irish Longitudinal Study on Ageing (TILDA), a nationally representative longitudinal cohort study of community-dwelling older adults in Ireland (Kearney et al., 2011; B. J. Whelan & Savva, 2013). From an initial 561 participants, 113 were excluded due to missing data, history of Parkinson’s disease, stroke, or transient ischaemic attack, presence of possible lesions, or fMRI data quality issues, and 154 were excluded for excessive head motion. Demographic information for both datasets is presented in Table 1.

**Table 1.**
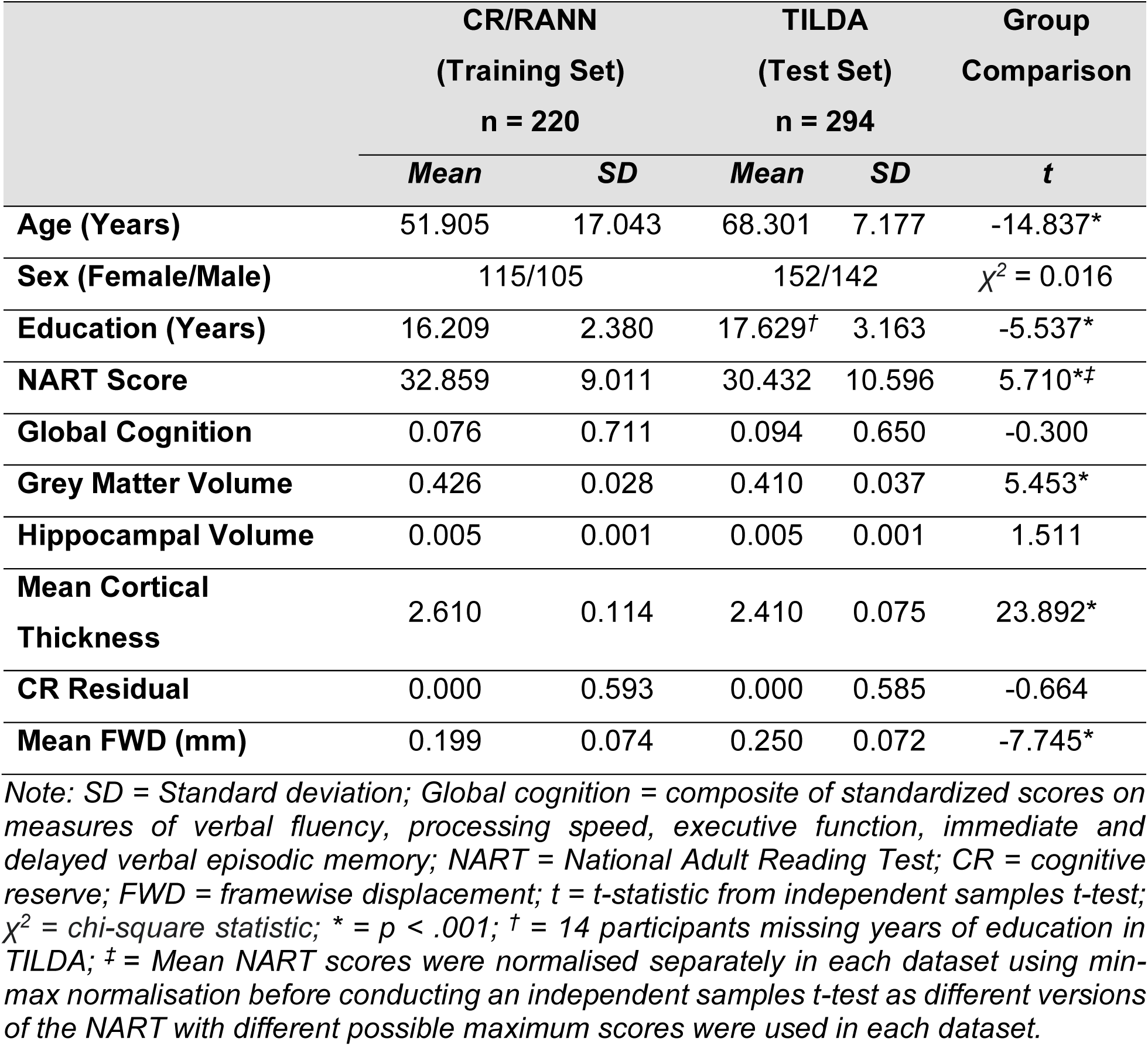
Descriptive statistics for relevant variables in both datasets.

|  | CR/RANN<br>(Training Set)<br>n = 220 |  | TILDA<br>(Test Set)<br>n = 294 |  | Group<br>Comparison |
| --- | --- | --- | --- | --- | --- |
|  | Mean | SD | Mean | SD | t |
| Age (Years) | 51.905 | 17.043 | 68.301 | 7.177 | -14.837* |
| Sex (Female/Male) | 115/105 | | 152/142 | | $\chi^2 = 0.016$ |
| Education (Years) | 16.209 | 2.380 | 17.629<sup>†</sup> | 3.163 | -5.537* |
| NART Score | 32.859 | 9.011 | 30.432 | 10.596 | 5.710*<sup>‡</sup> |
| Global Cognition | 0.076 | 0.711 | 0.094 | 0.650 | -0.300 |
| Grey Matter Volume | 0.426 | 0.028 | 0.410 | 0.037 | 5.453* |
| Hippocampal Volume | 0.005 | 0.001 | 0.005 | 0.001 | 1.511 |
| Mean Cortical<br>Thickness | 2.610 | 0.114 | 2.410 | 0.075 | 23.892* |
| CR Residual | 0.000 | 0.593 | 0.000 | 0.585 | -0.664 |
| Mean FWD (mm) | 0.199 | 0.074 | 0.250 | 0.072 | -7.745* |

Note: SD = Standard deviation; Global cognition = composite of standardized scores on measures of verbal fluency, processing speed, executive function, immediate and delayed verbal episodic memory; NART = National Adult Reading Test; CR = cognitive reserve; FWD = framewise displacement; t = t-statistic from independent samples t-test; $\chi^2$ = chi-square statistic; \* = $p < .001$ ; <sup>†</sup> = 14 participants missing years of education in TILDA; <sup>‡</sup> = Mean NART scores were normalised separately in each dataset using min-max normalisation before conducting an independent samples t-test as different versions of the NART with different possible maximum scores were used in each dataset.

### Image acquisition

#### Training Set

CR/RANN imaging data were obtained from a 3 T Philips Achieva scanner over the course of 2 separate 2-hour imaging sessions. Here, a single fMRI scan session was used, which was collected during completion of the Paper Folding task (Ekstrom et al., 1976), as described previously (Stern et al., 2014).The fMRI data were acquired using a 14 minute 26 second echo-planar imaging (EPI) pulse sequence (flip angle = 72°, slice thickness = 3 mm, slice gap = 0 mm, slices = 33, TR = 2000 ms, TE = 2 ms). In addition to 430 volumes, 3 dummy volumes were acquired at the start of the fMRI scan and automatically discarded. Structural MRI data were acquired using a 5-minute 3D T1- weighted magnetization-prepared rapid gradient echo (MPRAGE) scan with the following parameters: FOV = 256×256×180 mm, matrix size = 256×256, slice thickness = 1 mm, slice gap = 0 mm, TR = 6.5 ms, TE = 3 ms.

#### Training Set

TILDA imaging data were obtained using a 3 T Philips Achieva scanner during a 45-min MRI battery. Rs-fMRI data were acquired using a 6 minute 51.9 second gradient EPI sequence (flip angle = 90°, slice thickness = 3.2 mm, slice gap = 0.3 mm, slices = 38, TR = 2000 ms, TE = 28 ms). In addition to 200 volumes, 4 dummy volumes were acquired at the start of the rs-fMRI scan and automatically discarded. Structural MRI data were acquired using a 3D T1 MPRAGE scan with the following parameters: FOV = 240×240×162mm^3^, matrix size = 288×288, slice thickness = 0.9 mm, slice gap = 0 mm, TR = 6.7 ms, TE = 3.1 ms.

### Image Preprocessing

Each dataset was preprocessed separately with the same pipeline. Functional and structural images were manually reoriented to ensure approximately similar orientation in MNI space. Images were visually inspected for artefacts, data quality issues, possible lesions, and severe atrophy. Images were preprocessed using SPM12 and fMRI images were corrected for slice-timing and head motion. Nuisance regressors consisted of 6 motion estimates, mean WM signal, mean CSF signal, and mean global signal, and the derivatives, quadratic terms, and squares of derivatives of these 9 parameters (i.e., the ‘36 Parameter model’: (Ciric et al., 2017; Satterthwaite et al., 2013). Normalised functional images and variance images were visually inspected for data quality issues (e.g. registration or normalization errors) and motion-related issues and artefacts, respectively (*see Supplemental Information, Methods: Participant exclusions).* Finally, data were temporally smoothed with a zero-mean unit-variance Gaussian filter (approximate cut-off frequency of 9.37 Hz) using BioImageSuite (Joshi et al., 2011). The code used for quality control of the fMRI images is available here: https://github.com/rorytboyle/fMRI_QC.

### Functional connectivity network construction

The Shen 268-node functional atlas (Shen et al., 2013) was used to parcellate the fMRI data in both datasets, in line with previous CPM studies (Finn et al., 2015; M. Gao et al., 2020; Greene et al., 2018; Horien et al., 2019). Fully preprocessed functional volumes, already in MNI space, were resliced to the Shen functional parcellation image using spm_reslice. Using BioImageSuite, the mean time series for each node was calculated as the average time series across all voxels within each node, for each participant. Due to incomplete coverage of the cerebellum for a large proportion of the training set (n = 125; 56.82% of final sample), 63 nodes within the cerebellum and brainstem were removed from all participants in each dataset (*see Supplemental Information, Methods: Removal of cerebellar and brainstem nodes*). Functional connectivity between each pair of nodes was calculated by correlating the average time course between each pair of nodes. Pearson correlation coefficients were normalised by a Fisher z-transformation. This resulted in a 205 * 205 connectivity matrix for each participant in both datasets. The code used for construction of the connectivity matrices is available here: https://github.com/rorytboyle/fMRI_connectivity_processing.

### Measures

#### CR residual

CR residuals were obtained, separately in each dataset, from a linear regression of global cognition on age, gender, grey matter volume, hippocampal volume, and mean cortical thickness (see Fig. 1 and *Supplemental Information, Results: Creation of CR residuals*). Global cognition was measured as the composite of 5 standardized scores on comparable measures of verbal fluency, processing speed, executive function, immediate and delayed verbal episodic memory (*see Supplemental Information, Methods: Cognitive function and brain structure measures*). Measures of total GM volume, adjusted hippocampal volume, and mean cortical thickness were obtained from Freesurfer, as described previously (Carey et al., 2019; Habeck et al., 2016).

**Figure 1.**
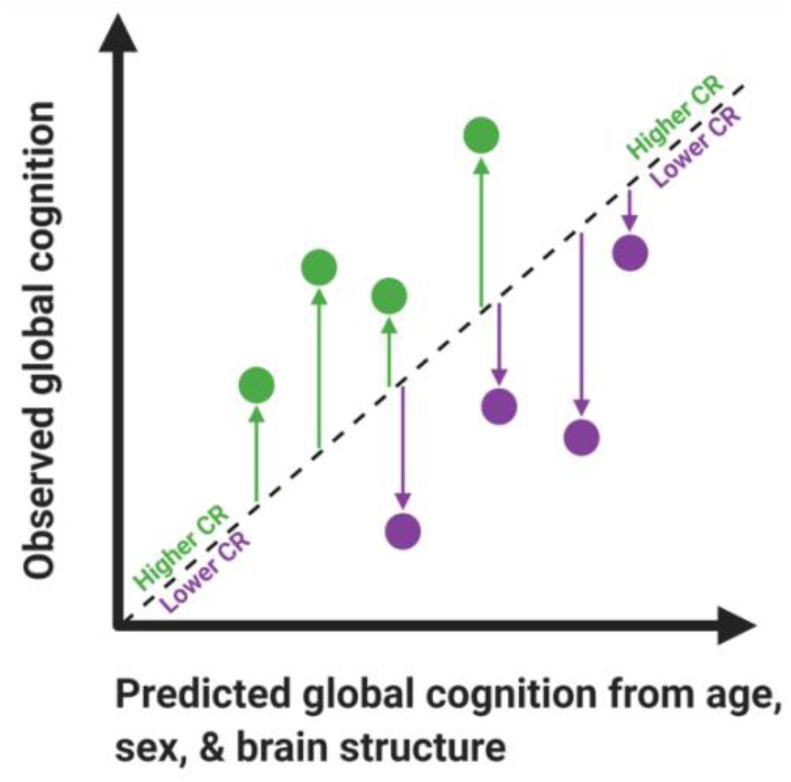
Illustration of CR residuals from the regression of global cognition on age, sex, and brain structure. Positive residuals (green arrows) reflect better cognitive performance than expected given age, sex, and brain structure. Negative residuals (purple arrows) reflect poorer cognitive performance than expected. Higher/more positive residual values reflect higher CR. Image adapted with permission from Fig. 1 (Franzmeier, Hartmann, et al., 2017).

#### CR proxy

Verbal intelligence was used here as CR proxy to assess the face validity of the neuroimaging measure of CR, as it is a robust socio-behavioural proxy of CR (Boyle et al., 2021). Verbal intelligence was measured by the total number of correctly pronounced words on the American National Adult Reading Test (AMNART; Grober & Sliwinski, 1991) in CR/RANN and on an adjusted version of the National Adult Reading Test (NART; Nelson & Willinson, 1982) in TILDA (*see Supplemental Information, Methods: Verbal intelligence)*.

### Connectome-based predictive modelling of cognitive reserve

CPM with leave-one-out cross-validation (LOOCV) was applied to the training set (Shen et al., 2017) using MATLAB (code available here: https://github.com/rorytboyle/flexible_cpm). CPM consisted of the following steps: edge selection, network strength calculation, model fitting, model application, model evaluation as detailed in Fig. 2 (*see Supplemental Information: Connectome-based predictive modeling* for a comprehensive description). This generated 3 network strength predicted CR values per participant *(positive network strength predicted CR, negative network strength predicted CR, and combined network strength predicted CR)*. The accuracy of each predicted value with respect to the CR residual was evaluated using 3 metrics: Pearson’s correlation, coefficient of determination (R^2^) from a linear regression, and the mean absolute error (MAE), in line with best-practice guidelines for predictive modelling in neuroimaging (Poldrack et al., 2020).

**Figure 2.**
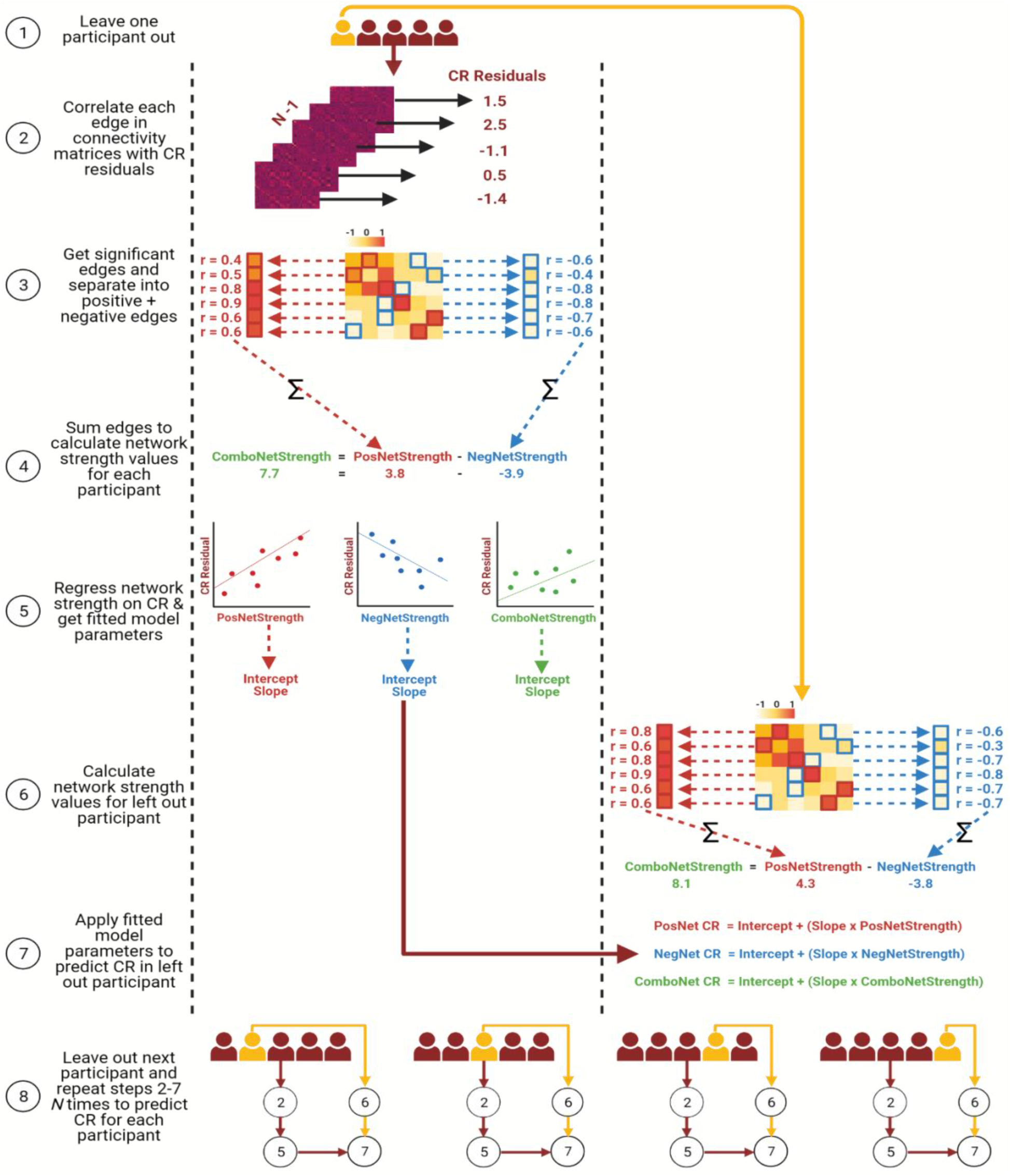
Schematic of CPM with LOOCV to predict CR residuals in the training set. PosNetStrength = positive network strength; NegNetStrength = negative network strength; ComboNetStrength = combined network strength; PosNet CR = positive network strength predicted CR; NegNet CR = negative network strength predicted CR; ComboNet CR = combined network strength predicted CR.

LOOCV is the standard cross-validation scheme in studies applying CPM (Greene et al., 2018; Rosenberg et al., 2016), but can overestimate model accuracy and can generate more variable predictions when applied to external datasets (Dwyer et al., 2018; Varoquaux et al., 2017). As such, k-fold cross-validation has been recommended as a preferable cross-validation scheme (Poldrack et al., 2020; Varoquaux et al., 2017). Therefore, the analysis was repeated in the training set using repeated k-fold cross-validation schemes (5-fold and 10-fold cross-validation repeated 100 times) instead of LOOCV (*see Supplemental Information: Repeated k-fold cross-validation)*.

### Optimisation of edge selection threshold

As edge-selection p-value thresholds are arbitrary (Greene et al., 2018), a data-driven method was implemented to obtain an optimal threshold which provided the highest training set accuracy (*see Supplemental Information, Methods: Optimisation of edge selection threshold).* The resulting p-value threshold was 0.0009 with r = 0.2896 (see Table S1 in *Supplemental Information, Methods*).

### Assessing validity of network strength predicted CR

To assess the theoretical validity of the network strength predicted CR measures, their face validity and protective effects on cognition were investigated. Face validity was assessed by establishing if there was a positive association with a CR proxy, verbal intelligence, using Pearson’s correlation. The protective effect was assessed by establishing whether they a) moderated the relationship between mean cortical thickness and global cognition (i.e., demonstrated a moderation effect), or b) were positively associated with global cognition, independent of mean cortical thickness (i.e., demonstrated an independent effect). In hierarchical linear regressions, global cognition was regressed on age, sex, and mean cortical thickness in Step 1, with network strength predicted CR added as an independent variable in Step 2, and the interaction term for mean cortical thickness and network strength predicted CR included as an independent variable in Step 3. The change in R^2^ (i.e., amount of variance explained) from Step 1 to Step 2, and from Step 2 to Step 3 in linear regression models were used to assess the size of the independent and moderation effects of CR proxies, respectively. This analysis was conducted in Python (code available here: https://github.com/rorytboyle/hierarchical_regression).

### External generalisability of connectome-based prediction

To evaluate if network strength predicted CR generalised to independent data, the trained CPM from CR/RANN was applied to TILDA data. First, positive and negative network strength values were computed by summing the positive and negative edges selected in each iteration of the LOOCV in the training set and dividing the sums by two to account for the symmetrical matrix. Combined network strength was computed as positive network strength minus negative network strength. Second, the regression parameters fitted in the training set were averaged across all iterations of the LOOCV and applied to their respective network strength values to calculate network strength predicted CR values. Third, these values were evaluated with respect to their predictive accuracy of the CR residual, using Pearson’s correlation, R^2^, and MAE. Finally, as described above for the training set, the predicted CR values were assessed with respect to their theoretical validity as measures of CR.

### Possible confounds in the relationship between connectivity and CR

The ’36 Parameter’ preprocessing pipeline used here has been shown to attenuate motion-related artifacts and noise in the data (Power et al., 2014; Yan et al., 2013). Due to the noted effect of motion on functional connectivity (Power et al., 2012), additional steps were also taken to control for this source of noise (*see Supplemental Information, Methods: Control of possible confounds).* To further ensure that the network strength predicted CR measures were not confounded by covariates including head motion, CPM was repeated including age, sex, and mean FWD as covariates at the edge selection step. This was implemented using a partial correlation to relate functional connectivity in each edge to the CR residual, including age, sex, and mean FWD as covariates.

## Results

### Connectome-based prediction of cognitive reserve

The connectome-based predictive models significantly predicted the CR residuals of novel participants (i.e., each left-out participant in the LOOCV) from task-based functional connectivity data in the training set (see Fig. 3 and Table 2). The combined network strength model had the highest predictive accuracy for the CR residual, across all 3 performance metrics (highest R, highest R^2^, and lowest MAE).

**Figure 3.**
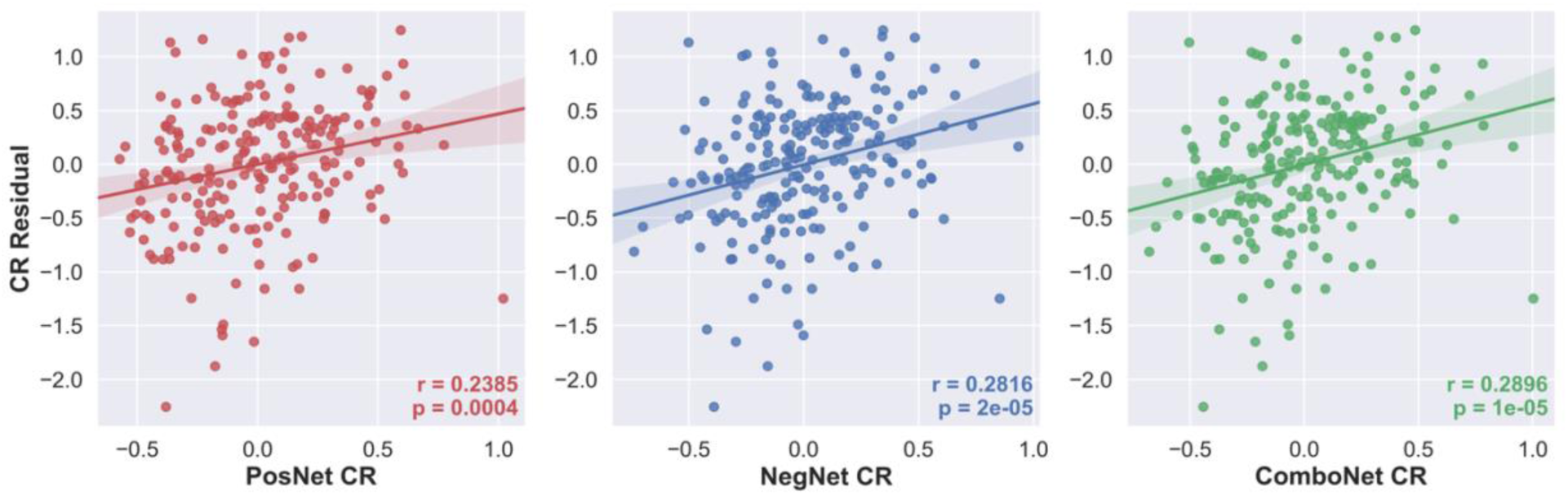
CR residual vs positive-, negative-, and combined network strength predicted CR in the training set.

**Table 2.**
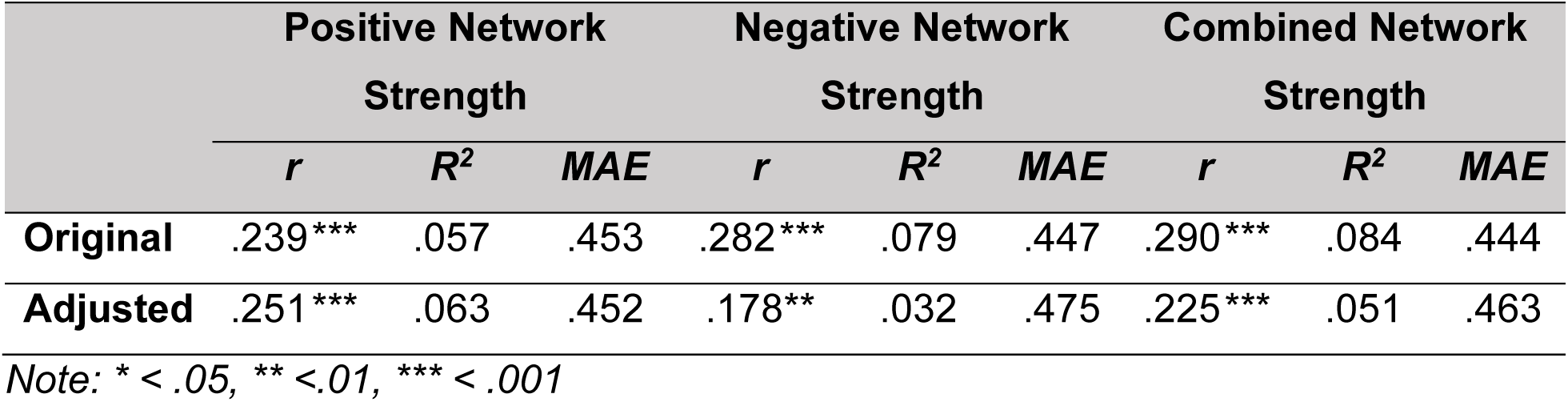
CPM performance for prediction of CR residuals in the training set.

### Validation of network strength predicted CR in the training set

The network strength predicted CR values generated by the connectome-based predictive models displayed face validity as measures of CR, as all models were significantly positively correlated with a CR proxy – verbal intelligence as measured by NART scores (see Fig. 4. and Table 3). The network strength predicted CR values also satisfied the protective effect criterion for measures of CR, as all were positively associated with global cognition, controlling for the effects of mean cortical thickness, age, and sex (see Table 3).

**Figure 4.**
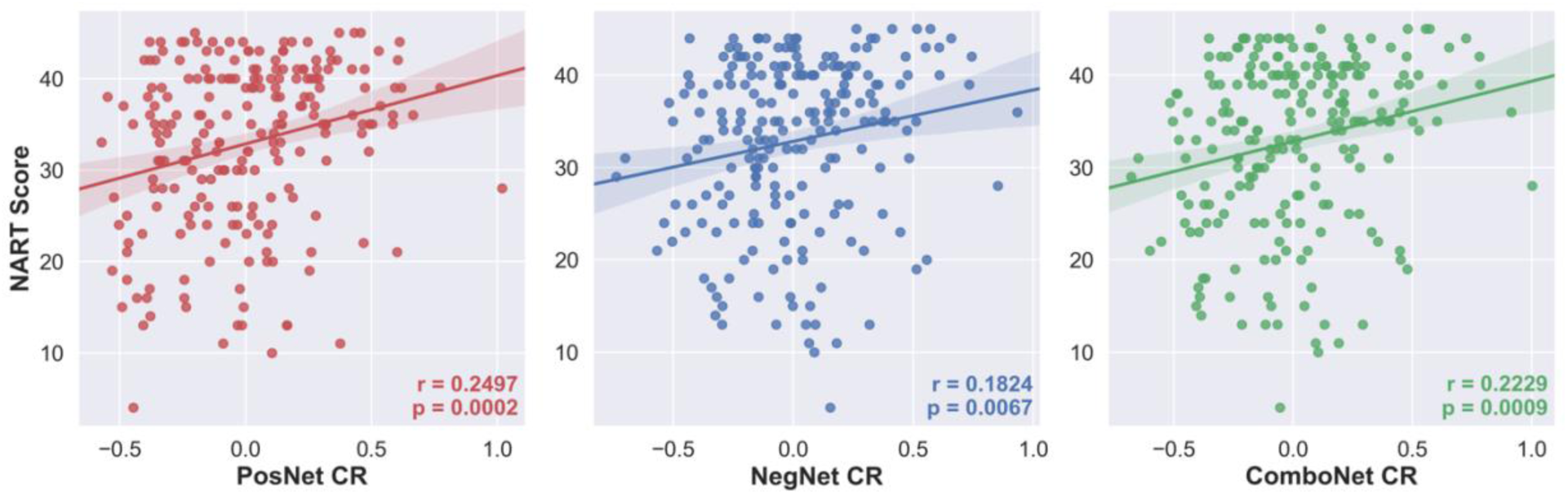
NART scores vs positive-, negative-, and combined network strength predicted CR in the training set.

**Table 3.** Validation of network strength predicted CR in the training set.

|  | Positive Network |  |  | Negative Network |  |  | Combined Network |  |  |
| --- | --- | --- | --- | --- | --- | --- | --- | --- | --- |
|  | Strength |  |  | Strength |  |  | Strength |  |  |
|  | NART | Ind. | Mod. | NART | Ind. | Mod. | NART | Ind. | Mod. |
|  | *r* | *R*<sup>2</sup> | *R*<sup>2</sup> | *r* | *R*<sup>2</sup> | *R*<sup>2</sup> | *r* | *R*<sup>2</sup> | *R*<sup>2</sup> |
| Original | .25*** | .043*** | 2.5e-5 | .182** | .06*** | .005 | .223*** | .063*** | .026 |
| Adjusted | .28*** | .03* | 1.7e-6 | .165* | .056*** | .003 | .226*** | .054*** | .002 |

Note: \* < .05, \*\* < .01, \*\*\* < .001. NART = Correlation of predicted values with NART scores; Ind. = Independent effect of predicted values on global cognition, controlling for age, sex, and mean cortical thickness; Mod. = Moderation effect of predicted values on relationship between brain structure and global cognition.

### Motion control and confounds

The connectome-based predictive models remained statistically significant when adjusting for age, sex, and mean FWD at the feature selection stage (see Fig. 5 and Table 2). Furthermore, network strength predicted CR values generated from the adjusted connectome-based predictive models also satisfied the criteria for measurement of CR as they displayed face validity and a positive independent effect on cognition (see Fig. 6 and Table 3).

**Figure 5.**
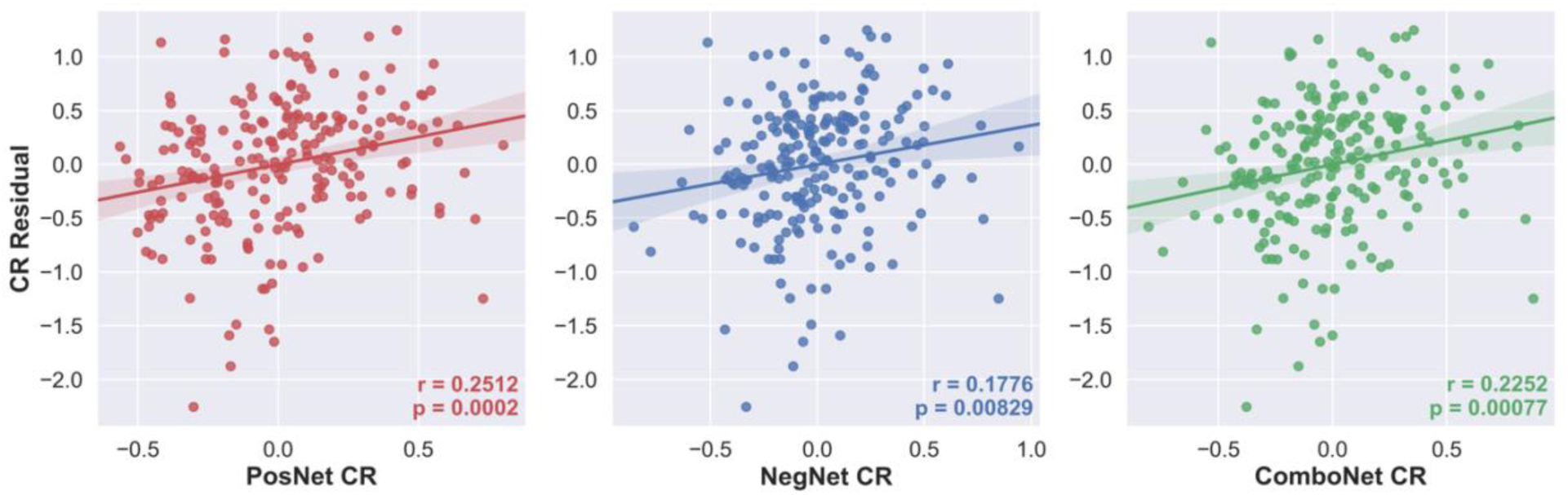
CR residual vs positive-, negative-, and combined network strength predicted CR using adjusted CPM in the training set.

**Figure 6.**
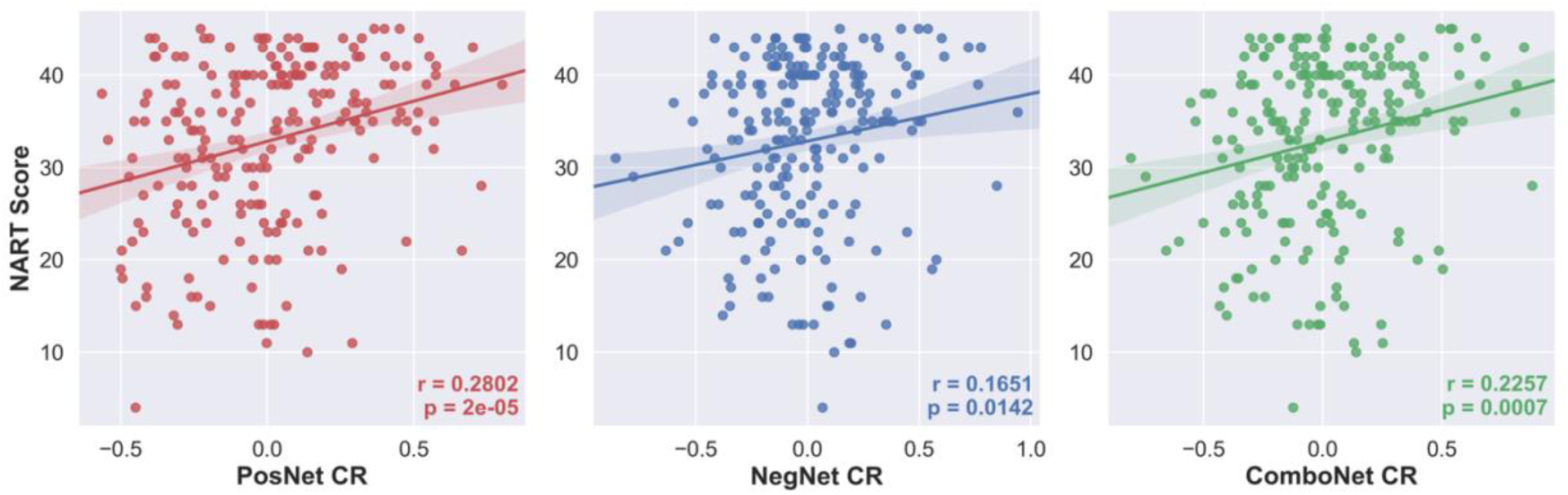
NART scores vs positive-, negative-, and combined network strength predicted CR using adjusted CPM in the training set.

### Functional network anatomy

Both positive and negative CR networks were sparse with 9 edges (0.04% of total edges) and 12 edges (0.06% of total edges) selected in every iteration of the positive and negative network, respectively (see Figs. 7 and 8). Nodes with multiple edges in the positive network were located within the left dorsolateral prefrontal cortex, left premotor/supplementary cortex, and the right angular gyrus (*see Table S4 in Supplemental Information, Results*). Nodes with multiple edges in the negative network were located in the left temporal pole, right angular gyrus and the left precentral gyrus (*see Table S5 in Supplemental Information, Results*).

**Figure 7.**
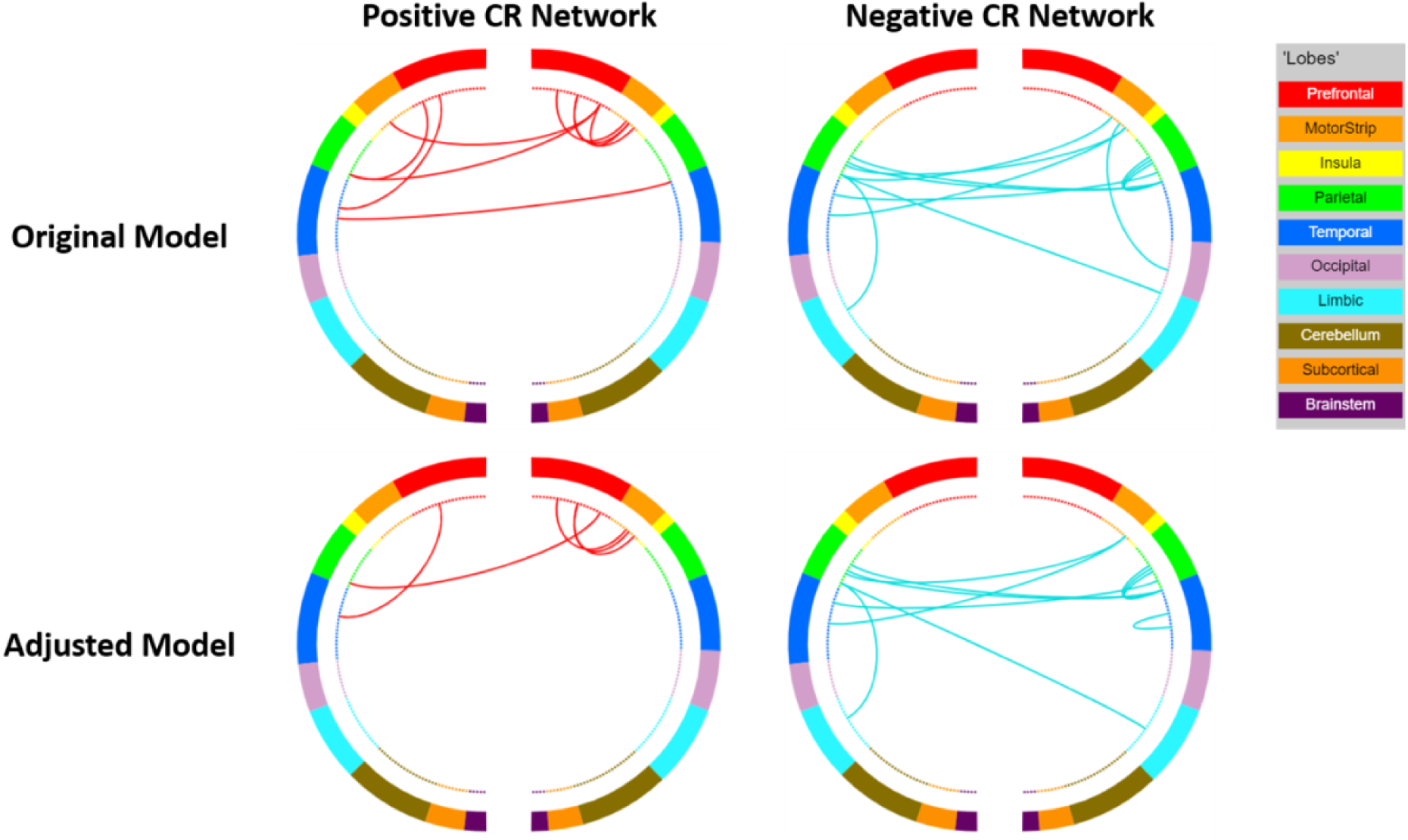
Circle plots illustrating the positive and negative CR connectomes. Positive connections (red) and negative connections (blue) in original CPM (top panel) and adjusted CPM (bottom panel), controlling for age, sex, and mean FWD. These circle plots are inverted such that the right side of each plot corresponds to the left hemisphere and the left side to the right hemisphere.

**Figure 8.**
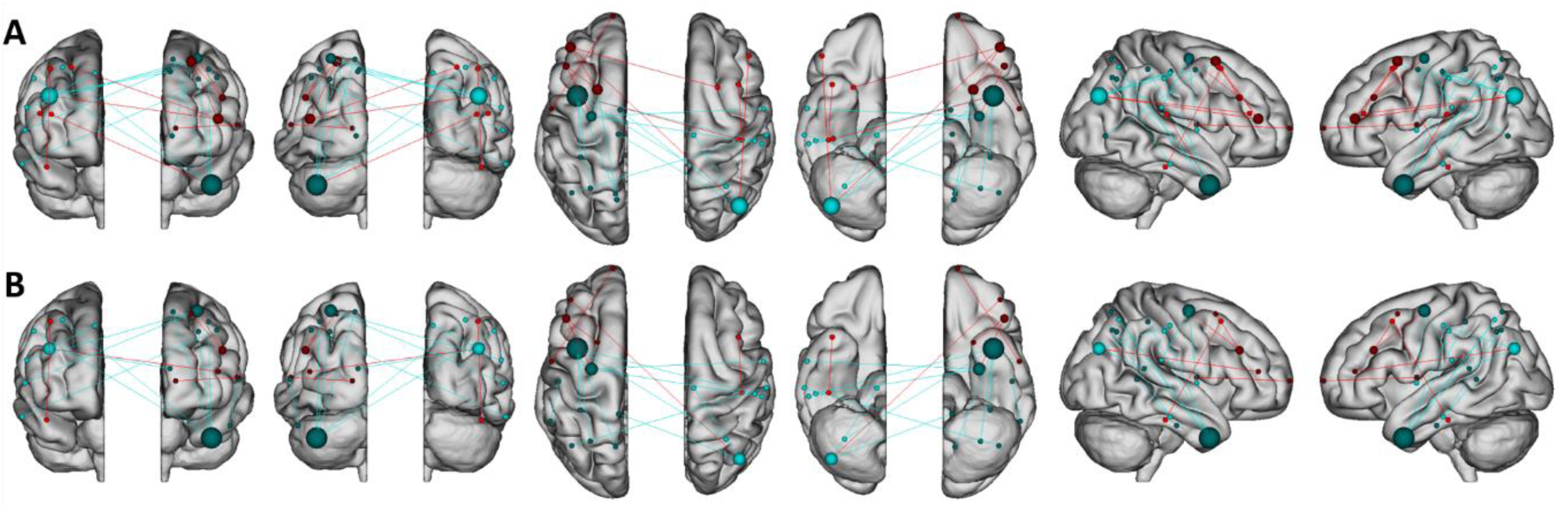
Glass brain visualising the patterns of connectivity within the brain. Positive connections (red) and negative connections (blue) in original CPM (A, top panel) and adjusted CPM (B, bottom panel), controlling for age, sex, and mean FWD.

In relation to canonical functional networks (Noble et al., 2017), the positive CR connectome was largely characterised by connectivity within the FPN, and of the FPN and motor network to other networks (see Fig. 9). The negative CR network was characterised by connectivity of a single medial frontal network node – the left temporal pole – to other networks, connectivity within the motor network, and connectivity of the motor network to other networks. Similar patterns of connectivity were observed in models adjusting for age, sex, and mean FWD, although a lower number of edges were selected.

**Figure 9.**
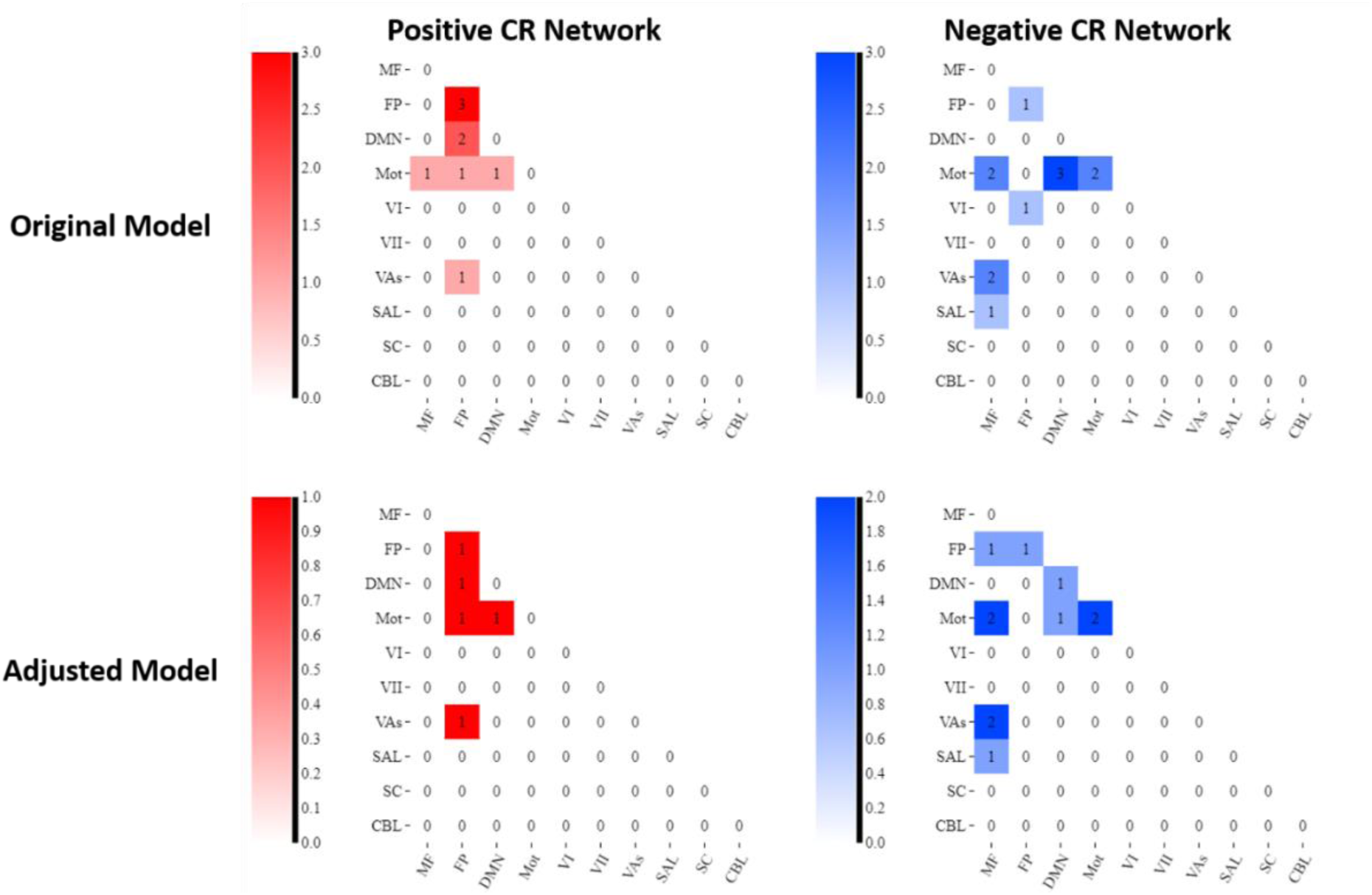
Connectivity matrices summarising the connectivity patterns within and between different functional networks. Note: Darker shades represent stronger connectivity (i.e., larger number of edges in that network). MF = Medial Frontal Network; FP = Frontoparietal Network; DMN = Default Mode Network; Mot = Motor Network; Vis I = Visual I Network; Vis II = Visual II Network; VAs = Visual Association Network; SAL = Salience Network; SC = Subcortical Network; CBL = Cerebellar Network.

### Generalisability of network strength predicted CR: Application to test set

The network strength predicted CR values in the test set were not related to the CR residual (see Fig. 10A and Table 4). While the correlations of the CR residual with negative- and combined-network strength predicted CR had p-values < 0.05, the negative direction of these associations meant that the associations are not meaningful as has been noted in other CPM studies (Ren et al., 2021). The results were similar when CPM, controlling for age, sex, and mean FWD, was applied to the test set (see Fig. 10B and Table 4).

**Figure 10.**
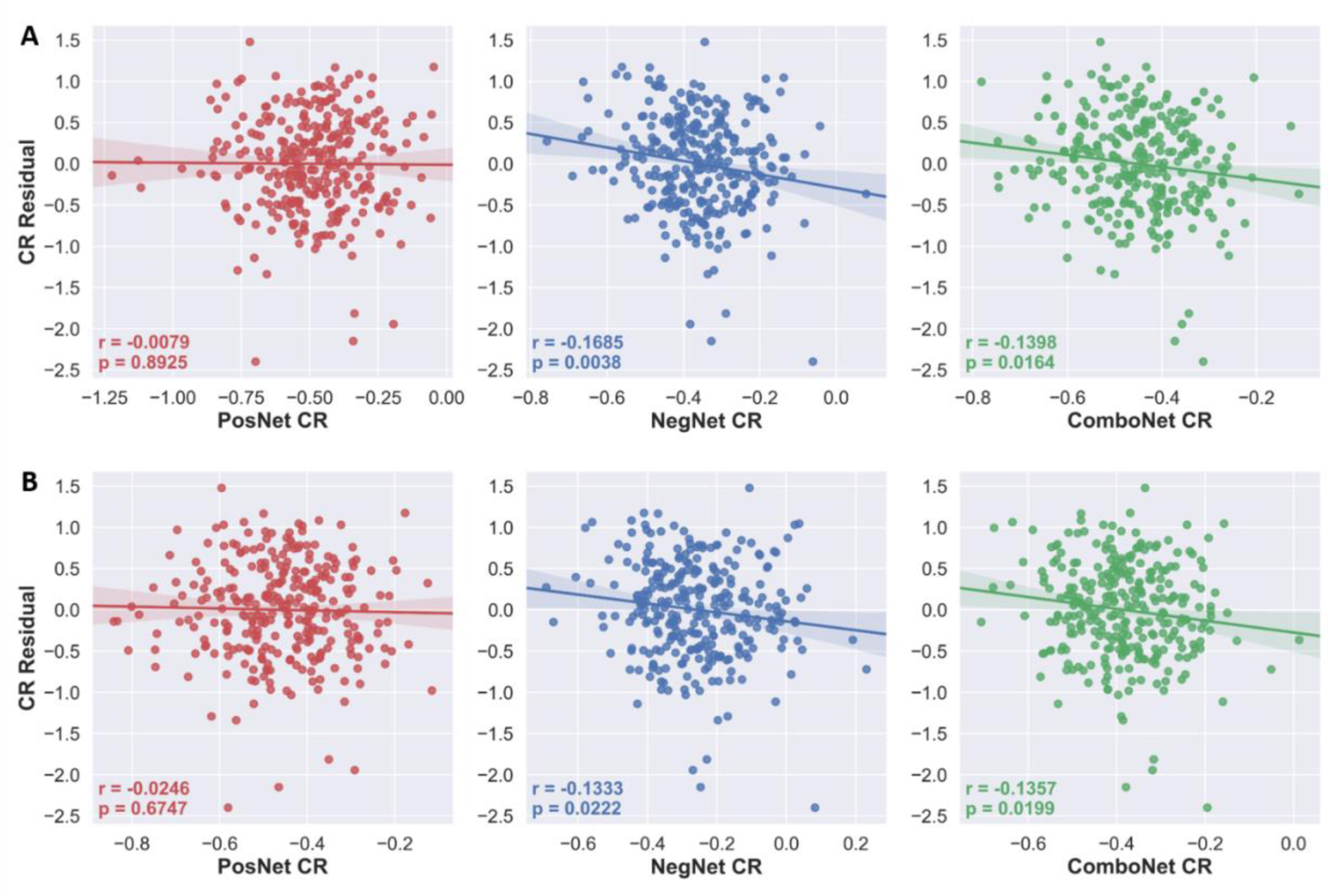
CR residual vs positive-, negative-, and combined network strength predicted CR in the test set. **A** = predicted CR values from original network strength models; **B** = predicted CR values from adjusted network strength models, controlling for age, sex, and mean FW.

**Table 4.**
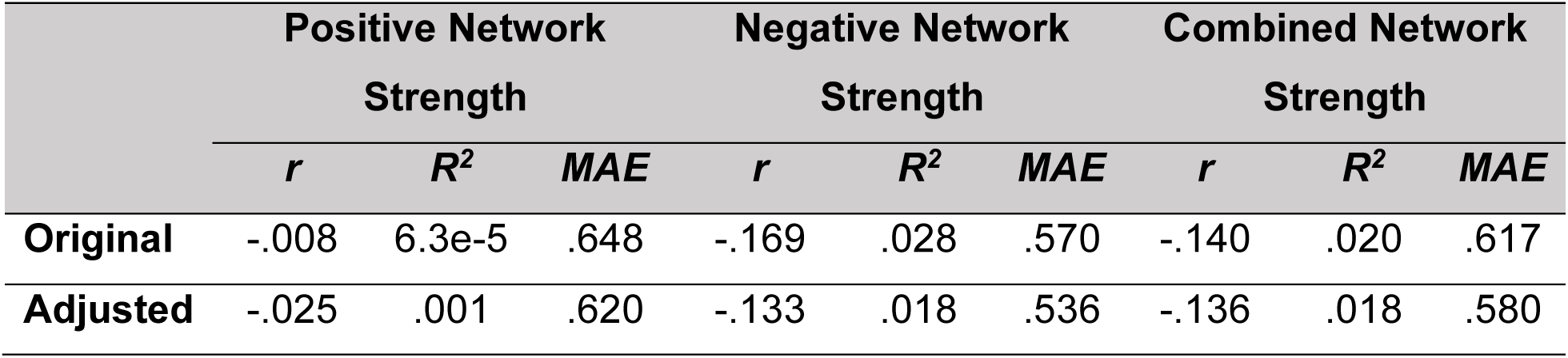
CPM performance for prediction of CR residuals in the test set.

|  | Positive Network |  |  | Negative Network |  |  | Combined Network |  |  |
| --- | --- | --- | --- | --- | --- | --- | --- | --- | --- |
|  | Strength |  |  | Strength |  |  | Strength |  |  |
|  | *r* | *R*<sup>2</sup> | *MAE* | *r* | *R*<sup>2</sup> | *MAE* | *r* | *R*<sup>2</sup> | *MAE* |
| **Original** | -.008 | 6.3e-5 | .648 | -.169 | .028 | .570 | -.140 | .020 | .617 |
| **Adjusted** | -.025 | .001 | .620 | -.133 | .018 | .536 | -.136 | .018 | .580 |

### Validation of network strength predicted CR in the test set

The network strength predicted CR values were not significantly positively correlated with a CR proxy, verbal intelligence (see Fig. 11A and Table 5). Furthermore, protective effects of the network strength predicted CR values on cognition were not observed as they did not moderate the relationship between mean cortical thickness and global cognition nor were they significantly positively associated with global cognition, controlling for the effects of mean cortical thickness, age, and sex (see Table 5). The adjusted models also did not demonstrate face validity (see Fig. 11B) or protective effects on cognition (see Table 5).

**Figure 11.**
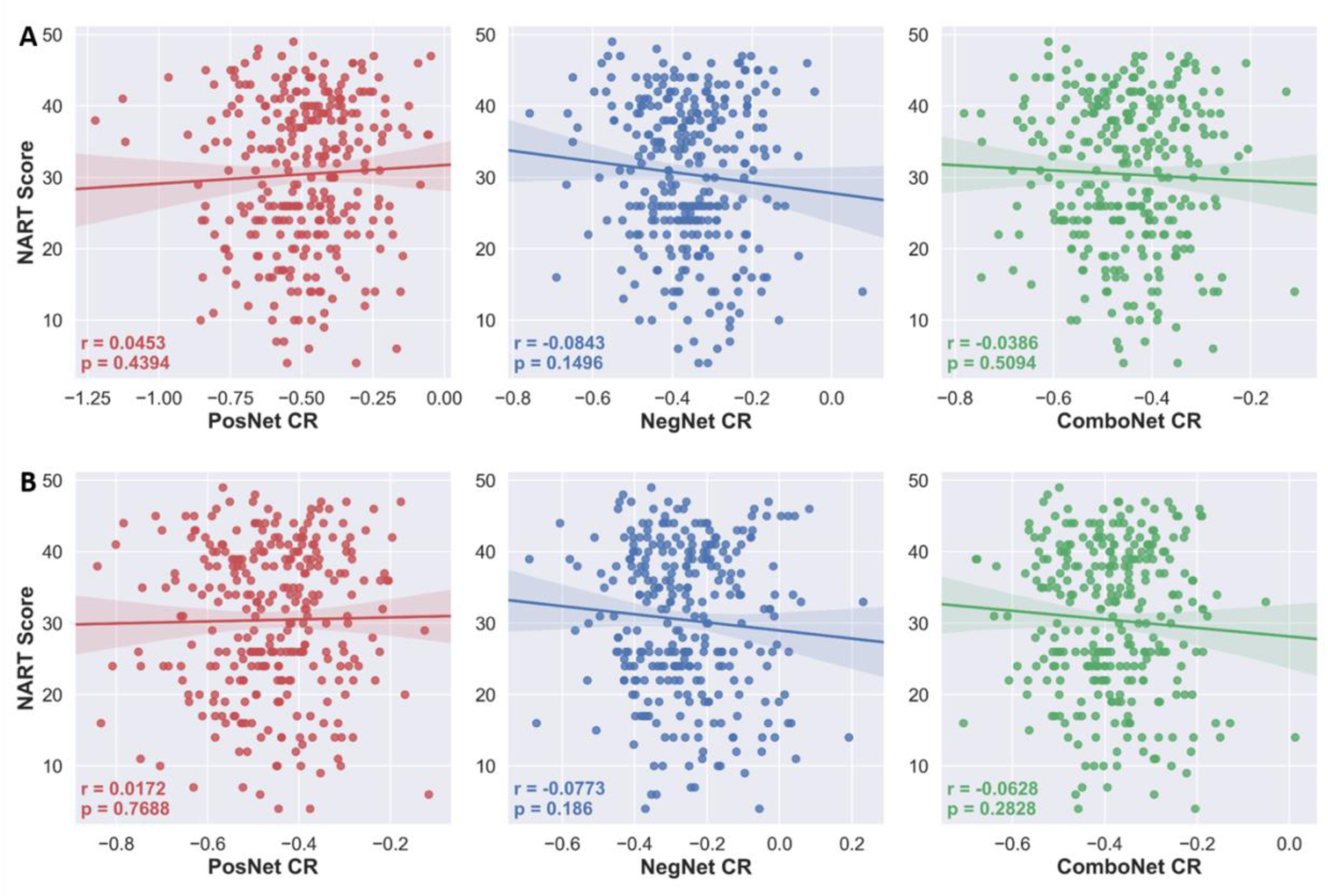
NART scores vs positive-, negative-, and combined network strength predicted CR in the test set. **A** = predicted CR values from original network strength models; **B** = predicted CR values from adjusted network strength models, controlling for age, sex, and mean FWD.

**Table 5.** Validation of network strength predicted CR in the test set.

|  | Positive Network |  |  | Negative Network |  |  | Combined Network |  |  |
| --- | --- | --- | --- | --- | --- | --- | --- | --- | --- |
|  | Strength |  |  | Strength |  |  | Strength |  |  |
|  | NART | Ind. | Mod. | NART | Ind. | Mod. | NART | Ind. | Mod. |
|  | *r* | *R*<sup>2</sup> | *R*<sup>2</sup> | *r* | *R*<sup>2</sup> | *R*<sup>2</sup> | *r* | *R*<sup>2</sup> | *R*<sup>2</sup> |
| **Original** | .045 | 2.9e-5 | .002 | -.084 | .024** | .001 | -.039 | .016* | .005 |
| **Adjusted** | .017 | 4e-4 | 4e-4 | -.077 | .015* | 6.1e-6 | -.063 | .015* | 3.5e-6 |

Note: \* < .05, \*\* < .01. NART = Correlation of predicted values with NART scores; Ind. = Independent effect of predicted values on global cognition, controlling for age, sex, and mean cortical thickness; Mod. = Moderation effect of predicted values on relationship between brain structure and global cognition.

### Post-hoc exploratory analyses

In the training set, the negative and combined network strength predicted CR values remained statistically significant when applying k-fold cross-validation methods instead of LOOCV (*see Table S6 in Supplemental Information, Results*). However, the positive network strength predicted CR values did not remain statistically significant in the training set. As in the main analysis, the k-fold models did not generalize to the test set as the negative- and combined-network strength predicted CR values were negatively correlated with the CR residual (*see Table S6 in Supplemental Information, Results*).

A negative correlation between network strength predicted CR values and observed values (i.e., the CR residual) is interpreted as a failure to explain any variance in the observed values (Greene et al., 2018) and is considered meaningless (Ren et al., 2021). To explore the negative correlation between network strength predicted CR values and the CR residual in the test set, the correlation between the CR residual and the thresholded edges in the test set was investigated (*see Table S7 in Supplemental Information, Results*). 12 edges were selected in the negative network in the training set. The average correlation between connectivity in these 12 edges and the CR residual in the training set was r = - 0.2728. However, in the test set, none of these edges were negatively correlated with the CR residual and the average correlation in the test set was r = 0.0696. Indeed, 3 of the thresholded edges (25%) were significantly positively correlated with the CR residual.

To investigate the failure of the model to generalize to rsfMRI data in an independent dataset, further exploratory analyses were conducted (*see Supplemental Information, Results: Exploratory Analyses)*. Applying CPM using a restricted age range in the training set to more closely match the age range of the test set failed to generalize to resting-state data in the test set (*see Table S8 in Supplemental Information, Results*). Applying CPM with a less conservative edge selection threshold (p = .01) also failed to generalize (*see Table S9 in Supplemental Information, Results*). Finally, using CPM to predict global cognition directly, controlling for measures of brain structure at the edge selection step, instead of predicting the CR residual, failed to generate measures that were positively associated with the CR proxy, verbal intelligence, in the training set.

## Discussion

CPM was applied to task-based functional connectivity to predict a CR residual in order to develop functional neuroimaging measures of CR, namely *positive-, negative-,* and *combined-network strength predicted CR.* These measures accurately predicted a CR residual in unseen individuals within the same dataset based on a sparse set of edges. The network strength predicted CR values met the theoretical criteria for neuroimaging measures of CR, as they displayed face validity and were positively associated with cognition beyond the effects of brain structure. However, these measures did not generalise to resting-state functional connectivity data from an independent dataset.

As was demonstrated in previous task-based fMRI studies, the network strength predicted CR measures here displayed face validity and protective effects on cognition. While previous studies used task-related activations (Stern et al., 2018) and task potency (van Loenhoud et al., 2020), the present study is the first to demonstrate that functional connectivity during task performance can predict CR in unseen data, albeit using internal cross-validation.

The CR connectomes identified here were sparse, reflecting connectivity strength from only 0.1% of all edges. This is more sparse than previously reported connectomes of cognitive phenotypes. For example, 3.88% and 1.99% of edges were included in connectomes underlying sustained attention (Rosenberg et al., 2016) and processing speed (M. Gao et al., 2020), respectively. The stricter edge selection threshold used here could explain this increased sparsity. This sparsity could also be a feature of CR-related connectomes, as similarly sparse functional networks have been reported to underlie CR. van Loenhoud et al. (2020) identified a sparse CR network comprising of 0.17% of all edges, although the same dataset as the present study was used, which does not rule out the possibility that the observed sparsity is an idiosyncrasy of the dataset.

The relationship between connectivity of the positive CR connectome and the CR residual in the present study was not robust despite being largely characterized by connectivity of the FPN which has been previously implicated in CR (Buckley et al., 2017; Franzmeier, Caballero, et al., 2017; Serra et al., 2016). Positive network strength predicted CR accurately predicted the CR residual in unseen data when CPM was implemented with LOOCV, but when implemented with k-fold cross-validation (*see Table S8 in Supplemental Information, Results*). LOOCV can produce estimates that have high variance (Efron, 1983), particularly compared to 10-fold cross-validation (Kohavi, 1995) and consequently can lead to overfitting (Lever et al., 2016). As such, the positive network strength predicted CR values may have somewhat reflected noise in the data (Poldrack et al., 2020; R. Whelan & Garavan, 2014).

Critically, the network strength predicted CR measures did not generalize to resting-state data in an independent dataset. Other studies applying CPM to cognitive phenotypes have had similar results, where the phenotype could be accurately predicted within-sample (i.e. in the training set), but not when applied to independent test sets (Gbadeyan et al., 2022; Manglani et al., 2021). We undertook exploratory analyses that suggested the failure to develop generalizable and theoretically valid measures of CR was not due to differences in the age ranges of the datasets, overfitting the training set due to a strict edge selection threshold, nor due to the use of the CR residual as a target variable. Another exploratory analysis investigated why the negative and combined network strength predicted CR values were negatively related to CR residuals in the test set. This was not a meaningful prediction because the predicted values were in the opposite direction to the observed values and similar findings have been treated as meaningless (Greene et al., 2018; Ren et al., 2021). Nevertheless, differences in connectivity from task to rest conditions may be particularly relevant to CR as adaptability of functional networks and processes to task demands is central to the CR construct. Inspection of the negative CR connectome in the test set revealed that all edges had positive, albeit mostly non-significant, correlations to CR. As these same edges were negatively correlated with CR in the training set, this may suggest that CR is associated with a change or reorganisation of brain connectivity in response to task demands, as previously shown by the relationship between task potency and CR (van Loenhoud et al., 2020). A practical implication of this is that while CR may be associated with both task-based and resting-state connectivity, the nature of these associations could be different. Therefore, it may not be possible for measures developed solely on task-based data to generalize to resting-state data, as has been demonstrated for CPM measures of cognitive phenotypes such as sustained attention (Rosenberg et al., 2016).

The inability to generalize to resting-state data may also have arisen due to the nature of the data in both our training and test set datasets. Single-task connectomes with static univariate functional connectivity were used in the training set but CPM studies have reported more accurate predictions of cognitive phenotypes with training sets consisting of multiple task connectomes (S. Gao et al., 2019), multivariate connectivity data (Yoo et al., 2019), and dynamic functional connectivity data (Zhu et al., 2021). In the test set, the resting-state fMRI scan was approximately five minutes in duration. This length is sufficient to obtain stable correlations for functional connectivity (Van Dijk et al., 2010), but longer durations further reduce the amount of noise in, and the reliability of, functional connectivity data (Birn et al., 2013; Van Dijk et al., 2010). The degree of individual variability in functional connectivity matrices is also greatly reduced in scans with fewer than 500 time points (Finn et al., 2015). As the test set resting-state scan contained only 200 time points, more time points may be needed for connectivity matrices to have sufficient variation across individuals in order to accurately predict complex phenotypes such as CR. Advanced modeling techniques, such as bootstrap aggregating (O’Connor et al., 2020) and partial least squares regression (Yoo et al., 2018), when implemented within CPM frameworks have also been shown to improve generalizability to external datasets.

There were some important limitations in the present study. Due to incomplete coverage of the cerebellum in a large proportion (57%) of the training set, nodes within the cerebellum and brainstem were removed from the functional connectivity matrices. This was necessary to avoid a drastic reduction in training set sample size but it reduced the number of edges in each connectivity matrix by 58%, from 35,778 to 20,910. The loss of this information in the training set, from a region that has been previously associated with CR (Belleville et al., 2021; Marques et al., 2016; Stern et al., 2018), may have hindered the ability to develop a generalizable measure of CR.

Other limitations arose due to the use of the CR residual as the target variable in the predictive model. The decision to use a CR residual instead of a CR proxy was justified on the basis that CR residuals are considered more direct measures of CR than proxy variables (Stern et al., 2020). However, because CR residuals inevitably contain a large amount of measurement error (Ewers, 2020), the use of the CR residual as a target variable, introduced irreducible error (i.e., noise in the dependent variable) into the predictive model. Irreducible error affects generalisability (Janssen et al., 2018), and can limit the ability of predictive models to reconstruct the target variables, due to the amount of noise present in that variable. In an exploratory analysis an alternative approach was implemented to minimize the effects of measurement error. This approach predicted cognition directly from functional connectivity, controlling for brain structure at the edge selection step of CPM. This approach also failed to generalize to the test set.

Despite the failure to generalize and the aforementioned limitations, there were a number of strengths to the current study. A data-driven approach was implemented that considered functional connectivity across the whole cortex. As such, the model and results were not biased by a priori predictions. A cross-validation framework was applied to assess whether the model could make accurate predictions in unseen data. An external validation dataset, with functional connectivity obtained from a different fMRI condition on which the model was trained, was used to provide a rigorous test of the generalisability of the developed measures across datasets and conditions. The gold-standard recommendations for deriving measures of CR (Collaboratory on Research Definitions for Reserve and Resilience in Cognitive Aging and Dementia, 2022) were rigorously applied by assessing the face validity of the measures in respect to their association with a robust socio-behavioural proxy of CR as well assessing their protective effects on cognition, above and beyond the effects of brain structure. Best-practice guidelines for predictive modeling in neuroimaging were also applied (Poldrack et al., 2020).

In sum, the present results demonstrated that task-based functional connectivity data can be used to create objective summary measures of CR (i.e., network strength predicted CR values) that are significantly associated with a CR residual, positively correlated with a CR proxy, and demonstrate a protective effect on cognition, beyond the effects of brain structure. These findings were demonstrated on unseen data within the training set (i.e., the same dataset used to develop the measures). However, the findings were not replicated when the model was applied to the test set (i.e., resting-state data from an independent dataset). The present study presents a framework for future attempts to develop measures that can generalise across datasets and fMRI conditions such that objective measures of CR can be developed, shared, and used by the wider research community with the ultimate aim of validating their clinical potential.

## Acknowledgements

The authors would like to thank all participants for the time and effort they generously gave to participate in the TILDA and CR/RANN studies. RB thanks Corey Horien, Stephanie Noble, Xenios Papademetris, Abigail Greene and Mehraveh Salehi for kindly and patiently responding to various queries about CPM methods, BioImageSuite, and anatomical labelling of the Shen atlas. RB also thanks Christian Habeck for helpful additional analysis ideas. MRI data collection in TILDA was supported by Prof. James Meaney and Dr. Jason McMorrow in the National Centre for Advanced Medical Imaging (CAMI) at St. James’ Hospital, Dublin.

This work was supported by grants to RB from the Irish Research Council (EPSPG/2017/277) and the NIH-supported Collaboratory on Research Definitions for Reserve and Resilience (10:GG013391-01). SPK is supported by Science Foundation Ireland (18/FRL/6188). YS is supported by NIA RF1 AG038465 and R01 AG026158. TILDA is funded by core grants from the Health Research Board, Atlantic Philanthropies and Irish Life. The TILDA MRI study was funded by the Health Research Board (HRA- PHR-2014-667). IHR thanks The Atlantic Philanthropies for their grant to the Global Brain Health Institute. The funding agencies had no involvement in the conduct of the research or preparation of the article.

Portions of the text in this manuscript were submitted as part of the first author’s PhD thesis. Figures 1 and 2 were made in BioRender (www.biorender.com) with publication license to RB.

