## Supplemental Information for "Connectome-based predictive modeling of cognitive reserve using task-based functional connectivity"

### Supplemental Methods

***Participant exclusions***

**CR/RANN:** From an initial dataset of 384 participants, participants were excluded according to the following criteria in the present study: (1) task fMRI data not available in session 1 or 2 of imaging data collection (3 participants excluded); (2) missing volumes in task fMRI scan (2 participants excluded); (3) presence of possible lesions or severe atrophy (5 participants excluded); (4) data quality issues including scanner artefacts, motion artefacts, and signal dropout (91 participants excluded); (5) excessive head motion defined as mean framewise displacement (FWD) > 0.4 mm or frame to frame movements > 97.5th percentile of head movements during resting-state fMRI scan (41 participants excluded); (6) missing data for CR residual (22 participants excluded).

**TILDA:** From an initial dataset of 561 participants, participants were excluded according to the following criteria: (1) structural and fMRI data available (5 participants excluded); (2) history of Parkinson’s disease, stroke, or transient ischaemic attack (11 participants excluded); (3) GM or WM lesion evident in structural MRI (19 participants excluded); (4) data quality issues including scanner artefacts, motion artefacts, and signal dropout (22 participants excluded); (5) excessive head motion defined as mean FWD > 0.4 mm or frame to frame movements > 97.5$^{th}$ percentile of head movements during resting-state fMRI scan (154 participants excluded); (6) missing data for CR residual (56 participants excluded)**.**

***Removal of cerebellar and brain stem nodes***

This approach was chosen to avoid excluding a large number of participants from the training set. Nodes within the cerebellum and brainstem were identified based on a previously reported anatomical labelling of the Shen parcellation atlas where the functional parcellations were assigned an anatomical label using the Tailarach atlas (Salehi et al., 2020). This resulted in the removal of 63 nodes in total, leaving a 430 (volume/time point) * 205 (node) and 200 (volume/time point) * 205 (node) time series for each participant in CR/RANN and TILDA, respectively.

***Connectome-based predictive modelling***

The implementation of LOOCV was such that this method iterated through the entire training set n times, with 1 participant left aside in each iteration for the model application step, and n-1 participants retained in each iteration for the edge selection, network strength calculation, and model fitting steps.

*Edge selection:* In each iteration, in the n-1 participants, each edge in the 205*205 connectivity matrix was correlated with the CR residual, using a Pearson's correlation. Edges with p-values below an optimised threshold of $p < .0009$ were selected (see *Optimising edge selection threshold)*. Thresholded edges were then separated into positive and negative edges, where positive edges were positively related to the CR residuals, and negative edges were negatively related to the CR residuals.

*Network strength calculation:* Positive and negative network strength values were then calculated for positive and negative edges separately. Positive Pearson's r values were summed and the sum was divided by 2 to account for matrix symmetry (i.e., the fact that each edge was represented twice in the symmetrical matrix). This was repeated for negative Pearson's r values. A combined network strength value was then calculated by subtracting negative network strength from positive network strength.

*Model fitting:* 3 separate linear regressions were fitted where positive/negative/combined network strength was the independent variable and the CR residual was the dependent variable. The model parameters (i.e., model intercept and regression coefficient/slope for network strength) were extracted from each regression.

*Model application*: Network strength values were then calculated for the left out participant. The edge strengths of the selected edges in the n-1 participants in the *Edge selection* step were summed for the left out participant. Summed edge strengths were then added to the fitted linear regression equations using the model parameters from the *Model fitting* step, in order to calculate 3 network strength predicted CR values (*positive network strength predicted CR, negative network strength predicted CR, combined network strength predicted CR)* for the left out participant.

For example:

PosNet CR = PosNet fitted intercept + (PosNet fitted slope * PosNet strength)

Where: PosNet = positive network; PosNet CR = positive network strength predicted CR; PosNet strength = summed edge strength of left out participant's positive network.

*Model evaluation:* In each iteration, the network strength predicted CR values were stored, as were the selected edges and the fitted model parameters. After n iterations, such that each participant was left out once, all participants had 3 network strength predicted CR values *(positive network strength predicted CR, negative network strength predicted CR, combined network strength predicted CR)*. Model performance, or the accuracy of each prediction, was evaluated for each network strength predicted CR value using 3 metrics: Pearson's correlation between network strength predicted CR values and the CR residual, coefficient of determination ($R^2$) from a linear regression with network strength predicted CR values as the independent variable and the CR residual as the dependent variable, and the mean absolute error (MAE) between the network strength predicted CR values and the CR residual.

***Repeated k-fold cross-validation***

In k-fold cross-validation, the data is randomly split into k subsets. One subset is then set aside (as the left out participant is set aside in LOOCV) as a test set for the *model application* step in CPM and the k-1 subsets are used to fit the model (i.e., *edge selection, network strength calculation, model fitting* steps in CPM). 100 iterations were run of each k-fold model, and the models were evaluated by averaging their Pearson's r and $R^2$ across datasets across the 100 iterations.

***Optimisation of edge selection threshold***

Starting with a p-value of 0.0001, CPM with LOOCV was repeated in the training set with 100 different p-value thresholds, increasing the p-value by 0.0001 at each step, to a maximum p-value of 0.01. The p-value threshold resulting in the largest Pearson's r between combined network strength predicted CR and the CR residual was selected as the optimal p-value threshold. This p-value threshold was 0.0009 with r = 0.2896.

**Table S1** Optimisation of edge selection threshold in the training set.

| p-value for Feature Selection | Positive Network Strength | | Negative Network Strength | | Combined Network Strength | |
|---|---|---|---|---|---|---|
| | r | p | r | p | r | p |
| 0.0001 | 0.1646 | 0.0145 | 0.2210 | 0.0010 | 0.2436 | 0.0003 |
| 0.0002 | 0.0560 | 0.4083 | 0.2022 | 0.0026 | 0.1895 | 0.0048 |
| 0.0003 | 0.0694 | 0.3058 | 0.1089 | 0.1072 | 0.1235 | 0.0675 |
| 0.0004 | 0.0335 | 0.6207 | 0.0915 | 0.1764 | 0.0846 | 0.2113 |
| 0.0005 | 0.0934 | 0.1676 | 0.1659 | 0.0137 | 0.1571 | 0.0197 |
| 0.0006 | 0.1304 | 0.0535 | 0.2116 | 0.0016 | 0.2022 | 0.0026 |
| 0.0007 | 0.1846 | 0.0060 | 0.2502 | 0.0002 | 0.2469 | 0.0002 |
| 0.0008 | 0.2324 | 0.0005 | 0.2687 | 0.0001 | 0.2783 | 0.0000 |
| 0.0009 | 0.2385 | 0.0004 | 0.2816 | 0.0000 | 0.2896 | 0.0000 |
| 0.0010 | 0.2152 | 0.0013 | 0.2849 | 0.0000 | 0.2818 | 0.0000 |
| 0.0011 | 0.1549 | 0.0215 | 0.2765 | 0.0000 | 0.2550 | 0.0001 |
| 0.0012 | 0.1337 | 0.0476 | 0.2744 | 0.0000 | 0.2464 | 0.0002 |
| 0.0013 | 0.0992 | 0.1423 | 0.2752 | 0.0000 | 0.2342 | 0.0005 |
| 0.0014 | 0.0728 | 0.2823 | 0.2751 | 0.0000 | 0.2238 | 0.0008 |
| 0.0015 | 0.0551 | 0.4161 | 0.2640 | 0.0001 | 0.2096 | 0.0018 |
| 0.0016 | 0.0445 | 0.5113 | 0.2577 | 0.0001 | 0.1990 | 0.0030 |
| 0.0017 | 0.0330 | 0.6265 | 0.2469 | 0.0002 | 0.1871 | 0.0054 |
| 0.0018 | 0.0344 | 0.6116 | 0.2504 | 0.0002 | 0.1879 | 0.0052 |
| 0.0019 | 0.0484 | 0.4751 | 0.2436 | 0.0003 | 0.1872 | 0.0053 |
| 0.0020 | 0.0606 | 0.3713 | 0.2559 | 0.0001 | 0.2006 | 0.0028 |
| 0.0021 | 0.0767 | 0.2575 | 0.2644 | 0.0001 | 0.2111 | 0.0016 |

| | | | | | | |
|---|---|---|---|---|---|---|
| 0.0022 | 0.0931 | 0.1686 | 0.2694 | 0.0001 | 0.2196 | 0.0010 |
| 0.0023 | 0.1071 | 0.1130 | 0.2705 | 0.0000 | 0.2235 | 0.0008 |
| 0.0024 | 0.1098 | 0.1042 | 0.2716 | 0.0000 | 0.2235 | 0.0008 |
| 0.0025 | 0.1045 | 0.1223 | 0.2731 | 0.0000 | 0.2212 | 0.0010 |
| 0.0026 | 0.1045 | 0.1221 | 0.2768 | 0.0000 | 0.2224 | 0.0009 |
| 0.0027 | 0.1005 | 0.1372 | 0.2756 | 0.0000 | 0.2194 | 0.0011 |
| 0.0028 | 0.0967 | 0.1531 | 0.2707 | 0.0000 | 0.2143 | 0.0014 |
| 0.0029 | 0.0988 | 0.1442 | 0.2729 | 0.0000 | 0.2139 | 0.0014 |
| 0.0030 | 0.0996 | 0.1408 | 0.2693 | 0.0001 | 0.2112 | 0.0016 |
| 0.0031 | 0.0869 | 0.1993 | 0.2666 | 0.0001 | 0.2046 | 0.0023 |
| 0.0032 | 0.0881 | 0.1931 | 0.2652 | 0.0001 | 0.2035 | 0.0024 |
| 0.0033 | 0.0949 | 0.1608 | 0.2611 | 0.0001 | 0.2034 | 0.0024 |
| 0.0034 | 0.0868 | 0.1997 | 0.2584 | 0.0001 | 0.1975 | 0.0033 |
| 0.0035 | 0.0945 | 0.1625 | 0.2600 | 0.0001 | 0.2011 | 0.0027 |
| 0.0036 | 0.0955 | 0.1581 | 0.2608 | 0.0001 | 0.2014 | 0.0027 |
| 0.0037 | 0.0910 | 0.1785 | 0.2603 | 0.0001 | 0.1989 | 0.0030 |
| 0.0038 | 0.0791 | 0.2427 | 0.2597 | 0.0001 | 0.1933 | 0.0040 |
| 0.0039 | 0.0791 | 0.2429 | 0.2560 | 0.0001 | 0.1906 | 0.0046 |
| 0.0040 | 0.0766 | 0.2580 | 0.2562 | 0.0001 | 0.1893 | 0.0048 |
| 0.0041 | 0.0777 | 0.2513 | 0.2575 | 0.0001 | 0.1892 | 0.0049 |
| 0.0042 | 0.0741 | 0.2738 | 0.2586 | 0.0001 | 0.1882 | 0.0051 |
| 0.0043 | 0.0703 | 0.2991 | 0.2590 | 0.0001 | 0.1875 | 0.0053 |
| 0.0044 | 0.0640 | 0.3448 | 0.2575 | 0.0001 | 0.1840 | 0.0062 |
| 0.0045 | 0.0632 | 0.3511 | 0.2574 | 0.0001 | 0.1831 | 0.0065 |
| 0.0046 | 0.0698 | 0.3024 | 0.2501 | 0.0002 | 0.1819 | 0.0068 |

| | | | | | | |
|---|---|---|---|---|---|---|
| 0.0047 | 0.0695 | 0.3047 | 0.2483 | 0.0002 | 0.1802 | 0.0074 |
| 0.0048 | 0.0730 | 0.2811 | 0.2454 | 0.0002 | 0.1791 | 0.0077 |
| 0.0049 | 0.0751 | 0.2674 | 0.2454 | 0.0002 | 0.1795 | 0.0076 |
| 0.0050 | 0.0761 | 0.2612 | 0.2385 | 0.0004 | 0.1758 | 0.0090 |
| 0.0051 | 0.0785 | 0.2461 | 0.2314 | 0.0005 | 0.1724 | 0.0104 |
| 0.0052 | 0.0798 | 0.2382 | 0.2219 | 0.0009 | 0.1669 | 0.0132 |
| 0.0053 | 0.0798 | 0.2384 | 0.2190 | 0.0011 | 0.1649 | 0.0143 |
| 0.0054 | 0.0812 | 0.2304 | 0.2183 | 0.0011 | 0.1654 | 0.0141 |
| 0.0055 | 0.0796 | 0.2398 | 0.2105 | 0.0017 | 0.1592 | 0.0182 |
| 0.0056 | 0.0800 | 0.2375 | 0.2070 | 0.0020 | 0.1572 | 0.0196 |
| 0.0057 | 0.0834 | 0.2180 | 0.2077 | 0.0020 | 0.1589 | 0.0183 |
| 0.0058 | 0.0871 | 0.1981 | 0.2057 | 0.0022 | 0.1594 | 0.0180 |
| 0.0059 | 0.0865 | 0.2011 | 0.2040 | 0.0024 | 0.1577 | 0.0193 |
| 0.0060 | 0.0871 | 0.1979 | 0.2014 | 0.0027 | 0.1563 | 0.0204 |
| 0.0061 | 0.0795 | 0.2403 | 0.1970 | 0.0033 | 0.1503 | 0.0258 |
| 0.0062 | 0.0852 | 0.2081 | 0.1927 | 0.0041 | 0.1504 | 0.0257 |
| 0.0063 | 0.0822 | 0.2244 | 0.1937 | 0.0039 | 0.1494 | 0.0267 |
| 0.0064 | 0.0812 | 0.2305 | 0.1923 | 0.0042 | 0.1476 | 0.0286 |
| 0.0065 | 0.0763 | 0.2600 | 0.1926 | 0.0041 | 0.1452 | 0.0313 |
| 0.0066 | 0.0728 | 0.2821 | 0.1843 | 0.0061 | 0.1390 | 0.0395 |
| 0.0067 | 0.0661 | 0.3289 | 0.1795 | 0.0076 | 0.1331 | 0.0486 |
| 0.0068 | 0.0629 | 0.3533 | 0.1806 | 0.0072 | 0.1324 | 0.0498 |
| 0.0069 | 0.0484 | 0.4749 | 0.1775 | 0.0083 | 0.1236 | 0.0673 |
| 0.0070 | 0.0412 | 0.5436 | 0.1787 | 0.0079 | 0.1210 | 0.0733 |
| 0.0071 | 0.0348 | 0.6075 | 0.1706 | 0.0113 | 0.1136 | 0.0927 |

| | | | | | | |
|---|---|---|---|---|---|---|
| 0.0072 | 0.0272 | 0.6877 | 0.1667 | 0.0133 | 0.1075 | 0.1117 |
| 0.0073 | 0.0206 | 0.7615 | 0.1664 | 0.0135 | 0.1042 | 0.1233 |
| 0.0074 | 0.0166 | 0.8063 | 0.1672 | 0.0130 | 0.1027 | 0.1290 |
| 0.0075 | 0.0120 | 0.8597 | 0.1654 | 0.0140 | 0.0992 | 0.1426 |
| 0.0076 | 0.0046 | 0.9461 | 0.1682 | 0.0125 | 0.0971 | 0.1513 |
| 0.0077 | -0.0028 | 0.9668 | 0.1658 | 0.0138 | 0.0917 | 0.1753 |
| 0.0078 | -0.0103 | 0.8791 | 0.1662 | 0.0136 | 0.0884 | 0.1913 |
| 0.0079 | -0.0122 | 0.8576 | 0.1633 | 0.0153 | 0.0859 | 0.2046 |
| 0.0080 | -0.0140 | 0.8364 | 0.1628 | 0.0157 | 0.0844 | 0.2124 |
| 0.0081 | -0.0213 | 0.7535 | 0.1608 | 0.0170 | 0.0799 | 0.2378 |
| 0.0082 | -0.0232 | 0.7322 | 0.1575 | 0.0194 | 0.0767 | 0.2571 |
| 0.0083 | -0.0290 | 0.6688 | 0.1593 | 0.0180 | 0.0745 | 0.2710 |
| 0.0084 | -0.0278 | 0.6820 | 0.1612 | 0.0167 | 0.0761 | 0.2609 |
| 0.0085 | -0.0272 | 0.6883 | 0.1599 | 0.0176 | 0.0756 | 0.2643 |
| 0.0086 | -0.0316 | 0.6409 | 0.1593 | 0.0181 | 0.0729 | 0.2816 |
| 0.0087 | -0.0332 | 0.6246 | 0.1611 | 0.0168 | 0.0728 | 0.2824 |
| 0.0088 | -0.0352 | 0.6033 | 0.1594 | 0.0180 | 0.0713 | 0.2923 |
| 0.0089 | -0.0334 | 0.6224 | 0.1615 | 0.0165 | 0.0733 | 0.2792 |
| 0.0090 | -0.0326 | 0.6302 | 0.1636 | 0.0152 | 0.0751 | 0.2671 |
| 0.0091 | -0.0288 | 0.6712 | 0.1657 | 0.0139 | 0.0786 | 0.2455 |
| 0.0092 | -0.0265 | 0.6961 | 0.1673 | 0.0129 | 0.0810 | 0.2315 |
| 0.0093 | -0.0256 | 0.7055 | 0.1675 | 0.0129 | 0.0812 | 0.2302 |
| 0.0094 | -0.0198 | 0.7702 | 0.1672 | 0.0130 | 0.0840 | 0.2148 |
| 0.0095 | -0.0182 | 0.7888 | 0.1675 | 0.0128 | 0.0849 | 0.2097 |
| 0.0096 | -0.0163 | 0.8099 | 0.1661 | 0.0136 | 0.0850 | 0.2094 |

| | | | | | | |
|---|---|---|---|---|---|---|
| 0.0097 | -0.0130 | 0.8482 | 0.1632 | 0.0154 | 0.0849 | 0.2096 |
| 0.0098 | -0.0074 | 0.9132 | 0.1634 | 0.0153 | 0.0876 | 0.1953 |
| 0.0099 | -0.0004 | 0.9958 | 0.1654 | 0.0141 | 0.0922 | 0.1728 |
| 0.0100 | 0.0050 | 0.9407 | 0.1610 | 0.0168 | 0.0925 | 0.1717 |

#### *Cognitive function and brain structure measures*

##### Cognitive function

**Verbal Fluency** was assessed using the total score on the Animals test in both datasets, which measures the ability to spontaneously produce the name of animals (Strauss et al., 2006). These variables – and all cognitive function measures – were standardised and Winsorized.

**Processing Speed** was measured using time to complete the Trail Making Test A (TMT; (Reitan, 1955) CR/RANN, and the Colour Trails Task 1 (CTT; (D'Elia et al., 1996) in TILDA. The CTT is considered a cross-culturally valid form of the TMT (Strauss et al., 2006). Scores were reversed coded, such that higher scores reflected greater cognitive performance.

**Executive Function** was assessed using the using time to complete the TMT B in CR/RANN and the CTT 2 in TILDA.

**Episodic Memory (Immediate)** was measured in CR/RANN using the total recall score on the Selective Reminding Test (SRT; (Buschke & Fuld, 1974), calculated by summing the total number of words recalled from 6 trials of a 12-item word list (Strauss et al., 2006). In TILDA, immediate verbal episodic memory was assessed by the average number of words immediately recalled from 2 trials of a 10-item word list as used originally in the Health and Retirement Study (Wallace & Herzog, 1995).

**Episodic Memory (Delayed)** was measured in CR/RANN using the delayed recall score from the SRT which consisted of the number of words recalled from the 12-item word list after an approximate 15 minute delay. In TILDA, immediate verbal episodic memory was assessed by the average number of words recalled from 2 trials of a 10-item word list after a 20-25 minute delay.

**Global Cognition** was measured using a composite measure of all five cognitive variables in each dataset: verbal fluency, processing speed, executive function, verbal episodic memory (immediate) and verbal episodic memory (delayed). All variables were Winsorized and standardised prior to creation of the composite. The composite variable was then Winsorized and standardised itself.

**Brain structure**

T1 MRIs were inspected and processed in TILDA and CR/RANN using FreeSurfer v6.0 and v5.1 (Fischl, 2012), respectively, as described previously (Carey et al., 2019; Habeck et al., 2016). Total GM volume and hippocampal volume were obtained from Freesurfer and were divided by estimated total intracranial volume to adjust for head size. Brain images were parcellated using the Desikan Killiany atlas, with 34 cortical regions of interest (ROIs) per hemisphere (Desikan et al., 2006). The mean cortical thickness of each cortical ROI was calculated. Mean cortical thickness was calculated as the mean over cortical ROIs. All variables were standardised and Winsorized.

##### *Verbal intelligence*

In TILDA, a measure (Strauss et al., 2006) was implemented to prevent undue stress/anxiety and to save time whereby participants completed the second half of the NART only if they scored greater than 20 on the first half. A correction procedure was used whereby scores of 0-11 were retained as full scores, but scores of 12-20 in participants who did not complete the second half were corrected using a conversion table outlined by Beardsall and Brayne (1990).

##### *Control of possible confounds*

First, images were visually inspected before and after preprocessing and excluded if motion-related artifacts were present. Second, participants with mean FWD > 0.5mm were excluded after preprocessing. Third, remaining participants who had individual head movements during the scan > 97.5$^{th}$ percentile of individual head movements across all participants were excluded. To assess whether functional connectivity was then related to head motion, the correlation between mean FWD and global functional connectivity (the average of functional connectivity in the upper triangle of the connectivity matrix) was assessed in each dataset. While the correlation in CR/RANN was not significant (r = .03, p = 0.62), global functional connectivity in TILDA was significantly related to mean FWD (r = 0.23, p < .001). As such, a final FWD threshold was applied such that participants with mean FWD > 0.4mm (n = 52) were excluded. After removal of these participants, the correlation between global functional connectivity and mean FWD did not change (r = 0.23, p < .001). Although the mean FWD > 0.4mm threshold did not reduce the correlation between mean FWD and global functional connectivity, there were fewer edges correlated to mean FWD at this threshold (18% fewer correlated edges). This threshold was chosen as it removed the remaining participants with the highest average head motion in TILDA (i.e., n = 52 with FWD <= 0.5 mm and > 0.4 mm) but retained a large final N. While a more conservative threshold of FWD > 0.2 mm has been previously used (Gao et al., 2020), this would have resulted in the exclusion of a further 95 and 239 participants from the final CR/RANN and TILDA samples respectively.

To ensure that the network strength predicted CR measures were not confounded by head motion, additional checks were conducted at the analysis stage to assess whether the CR residual was correlated with mean FWD and whether the network strength predicted CR measures were correlated with mean FWD. The possible influence of other confounds, specifically age and sex, on functional connectivity were assessed by correlating age with global functional connectivity and assessing gender differences in global functional connectivity. In the training set, mean FWD was not significantly associated with the CR residual (*Pearson's r = -.08, p = 0.25*) but was negatively associated with all network strength predicted CR values (*Positive r = -0.15, p = 0.02; Negative r = -0.14, p = 0.04; Combined r = -0.16, p = 0.02*). In the test set, mean FWD was not significantly associated with the CR residual (*r = -0.03, p = 0.64*) or network strength predicted CR values (*Positive r = - r = 0.02, p = 0.69; Negative r = 0.04, p = 0.53; Combined r = 0.04, p = 0.45*).

Age was positively associated with global functional connectivity in the training set (*r = 0.14, p = 0.03*) and negatively associated with connectivity in the test set (*r = - 0.18,* *p = 0.002*). Mean global functional connectivity did not differ between males and females in the training set (*Independent samples* $t = -0.11$, $p = 0.27$) but there was a significant sex difference in global functional connectivity in the test set ($t = 2.8917$, $p = 0.004$) such that males (*mean = 0.01, SD = 0.006*) had higher mean global functional connectivity than females (*mean = 0.008, SD = 0.005*).

Given the associations between network strength predicted CR and mean FWD in the training set, age and global functional connectivity in both datasets, and the sex differences in global functional connectivity in the test set, these three variables (FWD, age, and sex) were considered confounds. As such, an adjusted connectome-based predictive model was applied where, age, sex, and mean FWD were included as covariates at the feature selection stage, using a partial correlation between functional connectivity in each edge and the CR residual. After this adjustment, mean FWD was no longer significantly associated with negative network strength predicted CR ($r = -0.07$, $p = 0.28$) or combined network strength predicted CR ($r = -0.10$, $p = 0.13$). While mean FWD was still significantly associated with positive network strength predicted CR ($r = -0.13$, $p = 0.05$), the strength of the association was reduced as compared to the original (unadjusted) model. In the test set, as was also the case prior to adjusting for possible confounds, mean FWD was not significantly associated with positive network strength predicted CR ($r = 0.05$, $p = 0.42$); negative network strength predicted CR ($r = 0.04$, $p = 0.52$); and combined network strength predicted CR ($r = 0.06$, $p = 0.32$).

### Supplemental Results

#### *Creation of CR residuals*

The CR residuals accounted for 30% and 19% of the variance in global cognition in CR/RANN and TILDA respectively (see Table S2). The CR residuals accounted for a mean additional 57.9% of the variance in global cognition across both datasets (see Table S3). As such, the CR residuals were a suitable target variable for CPM as they reflected a larger amount of the variance in global cognition that was not explained by brain structure or demographics. In both datasets, CR residuals were approximately normally distributed and were significantly positively correlated with NART scores (see Fig. S1). As such, CR residuals in both datasets displayed face validity as measures of CR.

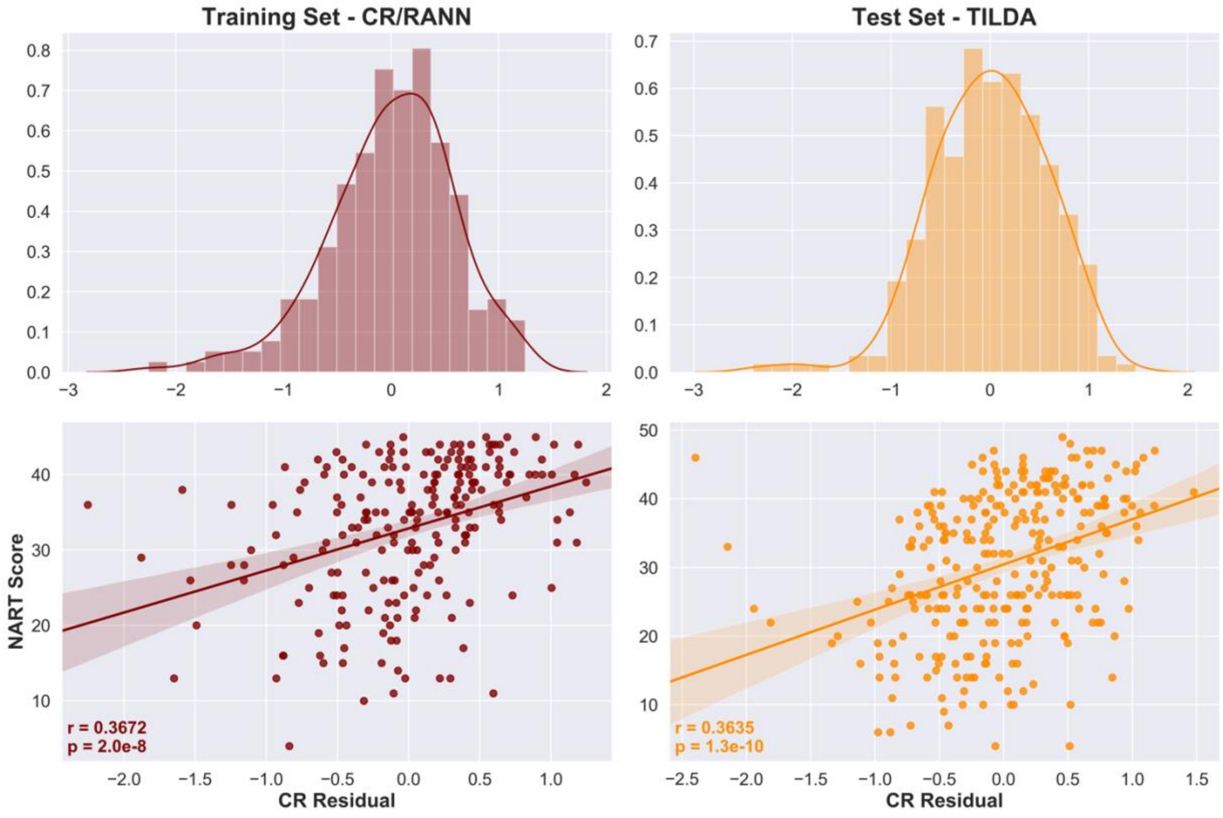


**Figure S1** Normality and face validity of CR residuals. *Histograms of CR residuals with kernel density estimates (top row) show that the CR residuals are approximately normally distributed. Scatterplots with regression lines (bottom row) show significant positive relationships between CR residuals and NART scores, demonstrating the face validity of CR residuals as CR measures.*

**Table S2** Results of multiple regressions used to create CR residuals in the training and test datasets.

| **Dataset** | **Model Statistics** | | | **GMV** | **HV** | **Cx Thick** | **Sex** | **Age** |
|---|---|---|---|---|---|---|---|---|
| | ***n*** | $R^2$ | ***f*** | ***β*** | ***β*** | ***β*** | ***β*** | ***β*** |
| CR/RANN (Training) | 220 | 0.304 | 18.723* | .088 | .001 | .080 | -.018 | -.433* |
| TILDA (Test) | 294 | 0.190 | 13.483* | -.045 | .077 | .040 | .095 | -.395* |

*Note: * = p < .0001, GMV = Grey Matter Volume, HV = Hippocampal Volume, Cx Thick = Mean Cortical Thickness*

**Table S3** Hierarchical regression results demonstrating average additional variance explained in global cognition across datasets.

| **Dataset** | **Step** | $R^2$ | $\Delta R^2$ | ***Additional % variance explained*** |
|---|---|---|---|---|
| **CR/RANN** | 1 | 0.293*** | | |
| | 2 | 0.988*** | .696 | 69.6% |
| **TILDA** | 1 | 0.188*** | | |
| | 2 | 0.651*** | .462 | 46.2% |
| **Average across datasets** | | | .579 | 57.9% |

*Note: *** = p < .001, Step 1 independent variables = age, sex, hippocampal volume; Step 2 independent variables = age, sex, hippocampal volume, CR residual*

#### *Functional network anatomy*

**Table S4**. Positive CR network functional neuroanatomy with nodes sorted by degree strength.

| # | BIS Label (BA) | Tailarach Label | K | Network | Lobe (L/R) | MNI co-ordinates | | |
|---|---|---|---|---|---|---|---|---|
| | | | | | | x | y | z |
| 154 | Dorsolateral prefrontal cortex (46) | Middle Frontal Gyrus | 3 | FP | L Prefrontal | -42.97 | 42.04 | 11.04 |
| 164 | Premotor/supplementary motor cortex (6) | Middle Frontal Gyrus | 2 | FP | L MotorStrip | -23.22 | 10.66 | 53.61 |
| 147 | Dorsolateral prefrontal cortex (9) | Middle Frontal Gyrus | 2 | FP | L Prefrontal | -46.12 | 28.15 | 26.79 |
| 49 | Angular Gyrus (39) | Middle Temporal Gyrus | 2 | DMN | R Parietal | 41.39 | -75.34 | 27.98 |
| 185 | Temporal Pole (38) | Superior Temporal Gyrus | 1 | MF | L Temporal | -38.01 | 6.07 | -37.86 |
| 166 | Premotor/supplementary motor cortex (6) | Precentral Gyrus | 1 | Mot | L MotorStrip | -27.58 | -9.08 | 55.86 |
| 163 | Premotor/supplementary motor cortex (6) | Superior Temporal Gyrus | 1 | Mot | L MotorStrip | -56.98 | -3.43 | 6.82 |
| 141 | Anterior Prefrontal Cortex (10) | Superior Frontal Gyrus | 1 | DMN | L Prefrontal | -11.7 | 65.09 | 4.18 |
| 62 | Primary Auditory Cortex (41) | Insula | 1 | Mot | R Temporal | 39.86 | -25.56 | 14.38 |

| | | | | | | | | |
|---|---|---|---|---|---|---|---|---|
| 59 | Inferior Temporal Gyrus (20) | Sub-gyral Temporal Lobe | 1 | VAs | R Temporal | 43.36 | -26.48 | -24.63 |
| 30 | Premotor/supplementary motor cortex (6) | Sub-gyral Frontal Lobe | 1 | FP | R MotorStrip | 25.22 | 12.41 | 49.39 |
| 19 | Dorsolateral prefrontal cortex (46) | Middle Frontal Gyrus | 1 | FP | R Prefrontal | 48.29 | 35.68 | 15.15 |
| 14 | Frontal Eye Fields (8) | Middle Frontal Gyrus | 1 | FP | R Prefrontal | 40.68 | 14.51 | 48.21 |

*Note: # = Node number; BIS Label (BA) = Brodmann Area label and number for node as listed in BioImageSuite; Tailarach Label = Anatomic label for node from Tailarach atlas* (Salehi et al., 2020)*; K = degree strength (i.e., number of connections) of node in positive network; Network = canonical networks defined in an independent sample* (Noble et al., 2017)*; Lobe (L/R) = Left or right hemisphere and Lobe as listed in BioImageSuite; MNI co-ordinates= Montreal Neurological Institute co-ordinates; FP = Frontoparietal Network; DMN = Default Mode Network; MF = Medial Frontal Network; Mot = Motor Network; VAs = Visual Association Network.*

**Table S5**. Negative CR network functional neuroanatomy with nodes sorted by degree strength.

| # | BIS Label (BA) | Tailarach Label | K | Network | Lobe (L/R) | MNI co-ordinates | | |
|---|---|---|---|---|---|---|---|---|
| | | | | | | x | y | z |
| 185 | Temporal Pole (38) | Superior Temporal Gyrus | 5 | MF | L Temporal | -38.01 | 6.07 | -37.86 |
| 49 | Angular Gyrus (39) | Middle Temporal gyrus | 3 | DMN | R Parietal | 41.39 | -75.34 | 27.98 |
| 166 | Premotor/supplementary motor cortex (6) | Precentral Gyrus | 2 | Mot | L MotorStrip | -27.58 | -9.08 | 55.86 |
| 164 | Premotor/supplementary motor cortex (6) | Middle Frontal Gyrus | 1 | FP | L MotorStrip | -23.22 | 10.66 | 53.61 |
| 218 | Premotor/supplementary motor cortex (6) | Cingulate Gyrus | 1 | Mot | L Limbic | -7.75 | -22.37 | 46.05 |
| 211 | Secondary Visual Cortex (18) | Lingual Gyrus | 1 | Vis I | L Occipital | -8.88 | -70.65 | -1.67 |
| 182 | Angular Gyrus (39) | Angular Gyrus | 1 | FP | L Parietal | -42.05 | -65.62 | 41.73 |
| 179 | Supramarginal Gyrus (40) | Sub-gyral Parietal Lobe | 1 | Mot | L Parietal | -35.72 | -39.37 | 47.75 |
| 178 | Visual Motor Co-ordination (7) | Precuneus | 1 | SAL | L Parietal | -9.83 | -66.34 | 55.14 |
| 177 | Visual Motor Co-ordination (7) | Sub-gyral Parietal Lobe | 1 | VAs | L Parietal | -28.41 | -62.35 | 40.42 |
| 161 | Premotor/supplementary motor cortex (6) | Cingulate Gyrus | 1 | Mot | L MotorStrip | -6.47 | -4.31 | 47.6 |

| | | | | | | | | |
|---|---|---|---|---|---|---|---|---|
| 89 | Dorsal Posterior Cingulate Cortex (31) | Cingulate Gyrus | 1 | Mot | R Limbic | 7.83 | -23.07 | 44.93 |
| 61 | Primary Auditory Cortex (41) | Superior Temporal Gyrus | 1 | Mot | R Temporal | 59.18 | -3.36 | 2.74 |
| 55 | Middle Temporal Gyrus (21) | Inferior Temporal Gyrus | 1 | FP | R Temporal | 61.28 | -22.87 | -22.38 |
| 46 | Supramarginal Gyrus (40) | Inferior Parietal Lobule | 1 | Mot | R Parietal | 58 | -29.28 | 19.53 |
| 45 | Supramarginal Gyrus (40) | Inferior Parietal Lobule | 1 | Mot | R Parietal | 52.84 | -27.25 | 40.93 |
| 43 | Visual Motor Co-ordination (7) | Precuneus | 1 | VAs | R Parietal | 31.62 | -60.79 | 49.21 |

*Note: # = Node number; BIS Label (BA) = Brodmann Area label and number for node as listed in BioImageSuite; Tailarach Label = Anatomic label for node from Tailarach atlas* (Salehi et al., 2020)*; K = degree strength (i.e., number of connections) of node in negative network; Network = canonical networks defined in an independent sample* (Noble et al., 2017)*; Lobe (L/R) = Left or right hemisphere and Lobe as listed in BioImageSuite; MNI co-ordinates= Montreal Neurological Institute co-ordinates; MF = Medial Frontal Network; DMN = Default Mode Network; Mot = Motor Network; FPN = Frontoparietal Network; Vis I = Visual I Network; SAL = Salience Network; VAs = Visual Association Network.*

##### *Exploratory Analyses*

First, CPM was repeated using a training set with a restricted age range of adults aged 50 years or older (n = 128, mean age = 64.42 yrs, SD = 8.49 years) to more closely reflect the age range of the test set. As in the main analysis, this model did not generalise to resting-state data in the test set (*see Table S8 in Supplemental Information, Results*). This suggested that differences in the age range in each dataset were responsible for the failure to generalize.

Second, CPM was repeated with a less conservative edge selection threshold of $p = .01$. With this threshold, negative network strength predicted CR values accurately predicted the CR residual, were significantly associated with verbal intelligence and demonstrated a protective effect on cognition (*see Table S9 in Supplemental Information, Results*). However, this model did not to generalize to the resting-state data in the test set. This suggested that the failure to generalize was not a result of inadvertent overfitting in the training set due to the data-driven method for optimization of the edge selection threshold selecting very few edges.

Third, CPM was repeated predicting global cognition instead of the CR residual, controlling for measures of brain structure (total GM volume, adjusted hippocampal volume, and mean cortical thickness) at the edge selection step. This was conducted to attenuate the influence of measurement error due to 1) the measurement error inherent in the CR residual itself and 2) the use of a two-step modelling process (i.e. linear regression to create CR residual, then CPM to predict CR residual). Global cognition was accurately predicted by negative and combined network strength predicted CR values (negative $r = .357$, combined $r = .31$, all $p < .01$), independent of brain structure, in the training set. However, these values were not significantly associated with the CR proxy, verbal intelligence, in the training set nor with verbal intelligence in the test set when applied to the rs-fMRI data. Therefore, this alternative model also failed to generate suitable neuroimaging measures of CR.

**Table S6** CPM performance for prediction of CR residuals in both datasets using k-fold cross-validation in the training set.

| CV Scheme | Dataset | Positive Network Strength | | Negative Network Strength | | Combined Network Strength | |
|---|---|---|---|---|---|---|---|
| | | ***r*** | ***$R^2$*** | ***r*** | ***$R^2$*** | ***r*** | ***$R^2$*** |
| **5-Fold** | Training | .087 | .010 | .197* | .041 | .175* | .032 |
| **5-Fold** | Test | -.021 | 4.2e-4 | -.139* | .019 | -.130* | .017 |
| **10-Fold** | Training | .093 | .011 | .212** | .046 | .187* | .036 |
| **10-Fold** | Test | -.079 | .006 | -.142* | .020 | -.164** | .027 |

*Note: * < .05, ** <.01*

**Table S7** Correlation of selected edges with CR residual in test set.

| Connectome | Node A | Node B | Networks | Training Set *r* | Test Set *r* |
|---|---|---|---|---|---|
| Positive | 59 | 14 | VAs – FP | 0.253** | 0.105 |
| Positive | 49 | 19 | DMN – FP | 0.261** | -0.037 |
| Positive | 154 | 30 | FP – FP | 0.248** | 0.022 |
| Positive | 154 | 49 | FP – DMN | 0.321*** | -0.021 |
| Positive | 185 | 62 | MF – Mot | 0.251** | 0.045 |
| Positive | 163 | 141 | Mot – DMN | 0.278*** | -0.085 |
| Positive | 164 | 147 | FP – FP | 0.237** | 0.016 |
| Positive | 166 | 147 | FP – FP | 0.241** | -0.075 |
| Positive | 164 | 154 | FP – FP | 0.237** | 2e-4 |
| Negative | 185 | 43 | FP – VAs | -0.277*** | 0.138* |
| Negative | 185 | 45 | FP – Mot | -0.315*** | 0.118* |
| Negative | 166 | 46 | Mot – Mot | -0.257** | 0.020 |
| Negative | 89 | 49 | Mot – DMN | -0.266** | 0.048 |
| Negative | 161 | 49 | Mot – DMN | -0.242** | 0.007 |
| Negative | 218 | 49 | Mot – DMN | -0.260** | 0.018 |
| Negative | 182 | 55 | FP – FP | -0.285*** | 0.071 |
| Negative | 166 | 61 | Mot – Mot | -0.265** | 0.034 |
| Negative | 211 | 164 | Vis I – FP | -0.248** | 0.106 |
| Negative | 185 | 177 | FP – FP | -0.309*** | 0.102 |
| Negative | 185 | 178 | FP – SAL | -0.269** | 0.121* |
| Negative | 185 | 179 | FP – Mot | -0.281*** | 0.053 |

*Note: * < .05, ** < .0005, *** < .0001. VAs = Visual Association Network; FP = Frontoparietal Network; DMN = Default Mode Network; MF = Medial Frontal Network; Mot = Motor Network; Vis I = Visual I Network; SAL = Salience Network.*

**Table S8** CPM performance for prediction of CR residuals in both datasets using an age-restricted sample in the training set.

| | Positive Network Strength | | | Negative Network Strength | | | Combined Network Strength | | |
|---|---|---|---|---|---|---|---|---|---|
| | ***r*** | ***$R^2$*** | ***MAE*** | ***r*** | ***$R^2$*** | ***MAE*** | ***r*** | ***$R^2$*** | ***MAE*** |
| **Original** | .142 | .020 | .563 | .209* | .044 | .527 | .195* | .038 | .536 |
| **Adjusted** | .079 | .006 | .707 | -.138* | .019 | .609 | -.076 | .006 | .680 |

*Note: * < .05*

**Table S9** CPM performance and validation of network strength measures in the training set using less conservative edge selection threshold.

| | Positive Network Strength | | | Negative Network Strength | | | Combined Network Strength | | |
|---|---|---|---|---|---|---|---|---|---|
| | ***Train*** | ***NART*** | ***Ind.*** | ***Train*** | ***NART*** | ***Ind.*** | ***Train*** | ***NART*** | ***Ind.*** |
| | ***r*** | ***r*** | ***rho*** | ***r*** | ***r*** | ***rho*** | ***r*** | ***r*** | ***rho*** |
| **P <.01 Adjusted** | -.001 | .147* | -.019 | .177** | .233*** | .227*** | .101 | .205** | .123 |

*Note: * < .05, ** < .01, *** <.001. rho = Partial Spearman's correlation between network strength predicted CR values and global cognition, controlling for brain structure.*
